# Optimal transport reveals unresolved cells in single-cell multi-omics

**DOI:** 10.64898/2026.10.15.759426

**Authors:** Mehran Piran, Fatemeh Vafaee

## Abstract

Single-cell multi-omics methods align cells across modalities, integrate them into one embedding and transfer labels between them. All three steps depend on a match between cells in two modalities, but this match usually stays hidden inside each tool. Its errors are hidden too, and every cell looks equally certain. Here we make the match explicit. For each pair of datasets, we build a soft coupling matrix with optimal transport and use it in three ways beyond estimating the match itself: to score how uncertain each cell’s match is, to upgrade existing integration and label-transfer tools without retraining them, and to decide which cells to set aside. Across 78 dataset pairs spanning six modality combinations, we evaluated ten integration and ten label-transfer tools using matched random removal as the control. We developed an ensemble coupling achieving the most accurate recovery of true cell pairs. Feeding couplings back into the tools improved label transfer clearly but improved integration only in how well the modalities mix. The ambiguity score, which we computed as a group-vote margin, gave the larger and more consistent gain in both tasks. Ambiguous-flagged cells matched worse in almost every dataset and removing them beat random removal in 98% of integration tests and 91% of label-transfer comparisons. These hard-to-integrate cells are mostly among rare, transitional and poorly measured populations. Making correspondence explicit therefore provides a practical way to distinguish reliable cross-modality results from uncertain ones while flagging cells that might have important roles in development and disease.

## Introduction

Single-cell technologies now measure several molecular layers of the same system, including transcriptomes, chromatin accessibility, surface proteins and DNA methylation, either jointly in the same cell (co-assays) or separately in comparable cell populations (1–4). These layers are informative only once they are aligned, which involves three related tasks: cross-modal alignment identifies which cells in one modality correspond to cells in the other, label transfer moves annotations from a reference to a query dataset, and data integration builds a shared representation in which similar cells from different assays sit close together. Although these tasks are usually handled and benchmarked separately (5–7), they all depend on the same underlying step: estimating correspondence between modalities. In most workflows, this correspondence remains hidden within the method and is neither independently assessed nor reused, with no cell-level measure of its uncertainty. Errors in this step can therefore affect both the integrated embedding and transferred labels, particularly when cross-modal signals are weak, sparse or indirect, as with ATAC-derived gene activity, targeted protein panels and methylation (5, 8).

This creates two main problems. First, hidden correspondence means hidden error: every cell in the returned embedding or label set looks equally certain, and benchmarks summarise a dataset with one number that does not say which cells carry the error. Second, many methods assign each cell to a single partner (5, 9). This is problematic in diagonal data, where no true cell-level counterpart exists, and in co-assay data, where noise may lead to incorrect hard matches. One-to-one matching also handles unequal cell numbers or compositions poorly and ignores alternative partners that may reveal uncertainty. Optimal transport (OT) offers a framework to reconnect unaligned single-cell data across time, modalities and perturbation conditions by estimating cell and population-level correspondences (10). In unpaired data, these represent plausible relationships between populations rather than hard cell-to-cell identity (9).

Optimal transport avoids hard one-to-one matching by allowing each cell to distribute its correspondence over several weighted partners. It finds the cheapest way to move the mass of one dataset onto the other, while fixed row and column sums prevent individual cells from becoming universal matches (9). We computed these couplings with entropic Sinkhorn regularisation, which enables efficient computations and produces smoother correspondence distributions (11). SCOT (12), SCOTv2 (13), Pamona (14) and uniPort (15) construct such couplings using Gromov-Wasserstein, partial or unbalanced transport, while UnionCom and MMD-MA align datasets without enforcing one-to-one matches (12, 13, 16). However, these methods use the correspondence mainly to construct an embedding and do not retain it for further analysis. Anchor-based methods (Seurat, Scanorama, and MNN with its faster implementation fastMNN) keep only sparse hard pairs (17–19), Harmony, Symphony, scVI and WNN hold no cell-level correspondence (20–23), and label-transfer tools (itClust, scmap, SingleR, Concerto) relate query to reference through centroids, correlations, neighbours or latent clusters (24–27). Other approaches such as scGLUE integrate unpaired modalities by combining modality-specific latent representations with prior regulatory relationships (28). Most of these tools therefore have no augmented approach. They report a prediction score, but that score lives inside one tool and one task and cannot be compared across tools or datasets, whereas a coupling gives every cell in both modalities a value on one common scale before any tool is applied. SONATA, the closest work to ours, detects geometric ambiguity in diagonal integration with Gromov-Wasserstein transport, but at the level of manifold regions rather than cells (8). To our knowledge, no existing method gives a tool-independent, per-cell measure of cross-modal match quality that works in co-assay and diagonal data and can be checked against a known pairing.

Here, we make cross-modal correspondence explicit by representing it as a soft optimal transport coupling matrix (Fig. 1). For each dataset pair, we use this coupling to address the four main aims of the study. The first aim is to estimate cross-modal correspondence and evaluate its accuracy. We developed four couplings based on different evidence: Diffusion-OT (local manifold shape), CCA-OT (shared feature variation), Diffusion-GW (within-modality geometry only) and Ensemble-OT (a consensus of Diffusion-OT, CCA-OT and the published MaxFuse method (29)). We compared them with SCOT, UnionCom and MMD-MA across 29 co-assay datasets, where true cell partners are known, using the fraction of samples closer than the true match (FOSCTTM). Although Ensemble-OT was most accurate overall, the best coupling varied by modality pair and downstream tool. We therefore retained Diffusion-OT, CCA-OT, Ensemble-OT and MaxFuse for later analyses, while using the less accurate Diffusion-GW only as a geometry-only comparison. The second aim is to quantify coupling uncertainty, which we call ambiguity. Ambiguity reflects uncertainty in the matching process rather than an intrinsic property of the cell. For each cell, we pool the coupling mass by groups in the opposite modality. A cell is considered ambiguous when the two most strongly supported groups receive similar coupling mass. LF-Amb (Label-free ambiguity detection method) uses unsupervised clusters for integration, whereas RefA-Amb (first modality as reference) and RefB-Amb (second modality as reference) use labelled reference cell types for label transfer in each direction. We then tested whether flagged cells were truly poorly matched and which dataset properties were associated with ambiguity.

**Figure 1.**
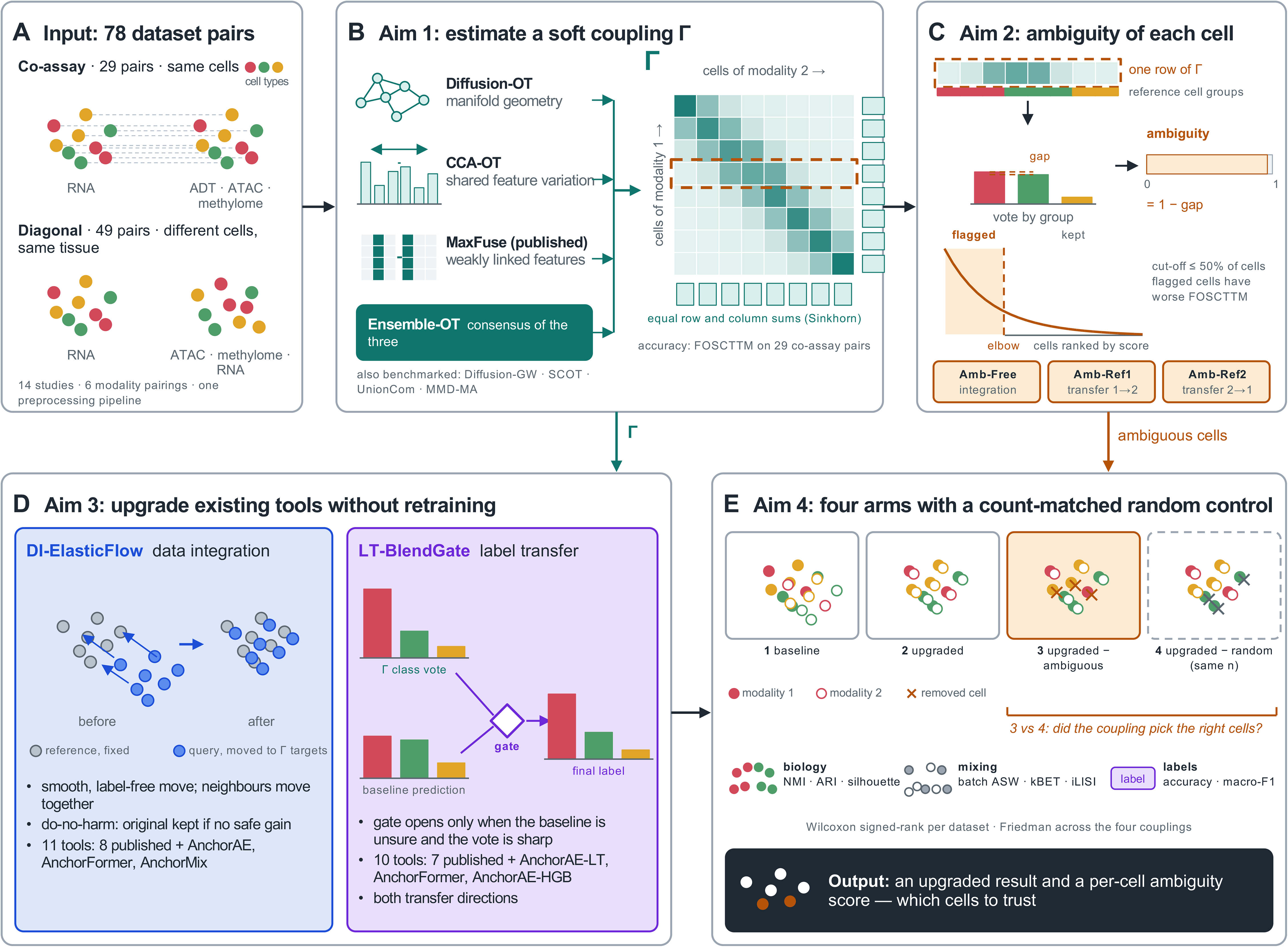
Overview of the framework. (A) Input. Each dataset pair has two modalities. In co-assay pairs both modalities are measured in the same cells, so the true partner of every cell is known (29 pairs). In diagonal pairs they are measured in different cells of the same tissue (49 pairs). The benchmark holds 78 pairs from 14 studies and six modality pairings, all processed with one pipeline. (B) Aim 1. Four couplings estimate the correspondence between the modalities: Diffusion-OT (manifold shape), CCA-OT (shared feature variation), the published tool MaxFuse (weakly linked features) and Ensemble-OT, a rank consensus of the three. Each yields a soft, non-negative coupling matrix Γ whose rows are cells of modality 1, columns are cells of modality 2 and entries are the correspondence mass shared by a cell pair; Sinkhorn balancing fixes the row and column sums. Accuracy is measured by FOSCTTM on the co-assay pairs. Diffusion-GW, SCOT, UnionCom and MMD-MA were benchmarked for accuracy only. The dashed row marks a cell whose mass is split between two groups of partners. (C) Aim 2. For each cell, coupling mass is pooled by cell group in the other modality, and ambiguity is one minus the gap between the two strongest votes. LF-Amb uses unsupervised clusters on both sides (data integration); RefA-Amb and RefB-Amb use the cell types of the labelled reference, modality 1 or modality 2 (label transfer in each direction). The cut-off is the elbow of the ranked scores, capped at 50% of cells. (D) Aim 3. DI-ElasticFlow keeps the reference embedding fixed and moves query cells smoothly toward their coupling targets, choosing candidates without cell-type labels and returning the original embedding if none passes the do-no-harm guard. LT-BlendGate pools coupling mass per reference cell type into a class vote and blends it with the baseline prediction through a gate that gives the coupling weight only when the baseline is unsure and the vote is sharp. Eleven integration tools and ten label-transfer tools were tested (Methods). (E) Aim 4. Every tool, coupling and dataset is evaluated in four arms; comparing arms 3 and 4 isolates the value of choosing which cells to remove. Metrics and tests are listed in the panel.

The third aim is to improve existing tools without retraining them by using the coupling as an independent source of evidence. We developed DI-ElasticFlow, which keeps reference cells fixed and moves query cells smoothly toward coupling-derived positions. A single label-free Safe-upgrade rule accepts a correction only when modality mixing improves without weakening local neighbourhood structure or cell-type separation; otherwise, it returns the original embedding. LT-BlendGate converts the coupling into a vote over reference cell types and blends it with the baseline prediction, giving the coupling more weight when the baseline is uncertain and the vote is clear. We applied these upgrades to eleven integration and ten label-transfer tools, including three integration models developed here: AnchorAE, AnchorFormer and AnchorMix. AnchorAE and AnchorFormer were also paired with label-transfer classifiers. This range was chosen because no single method suits every modality pair and our custom models test the upgrades across different architectures (30). The fourth aim is to determine whether identifying ambiguous cells improves analysis of the remaining cells. For each tool and coupling, we compared four conditions: the original result, the upgraded result, and the upgraded result after removing either ambiguous cells or an equal number of random cells. The matched random control determines whether improvement comes from selecting the right cells rather than simply removing cells. When the Safe-upgrade rule accepts no correction, removal is applied to the original embedding.

We evaluated the framework on 78 dataset pairs from fourteen studies, spanning six modality combinations in both co-assay and diagonal designs. Making cross-modal correspondence explicit through a soft coupling captured information beyond the integrated embedding. Incorporating this coupling substantially improved label transfer and enhanced cross-modal mixing in data integration. Its most consistent contribution was quantifying correspondence ambiguity across tasks, methods and modality pairs. Ambiguous cells were enriched near cell-type and modality boundaries, especially among rare, immature, transitional and poorly resolved populations. Removing these cells improved performance more than coupling-based correction in both transfer directions. Coupling choice had little effect on embedding correction but strongly affected ambiguity detection. Overall, explicit soft correspondence provides an interpretable, cell-level measure of cross-modal confidence that supports more transparent multi-omics analysis.

## Results

We assembled 78 dataset pairs from 14 published studies, covering six modality pairings in two regimes. The 29 co-assay pairs measured both modalities in the same cells (15 RNA–ADT, 12 RNA–ATAC and 2 RNA–methylome), whereas the 49 diagonal pairs measured them in different cells from the same tissue (25 RNA–ATAC, 6 RNA–methylome and 18 RNA–RNA). Dataset details, including cell counts, cell types, source references and accession identifiers, are provided in Datasets_composition_supplementary.pdf. Figure 1 summarizes the framework: input data (Fig. 1A), coupling inference (Fig. 1B), ambiguity scoring (Fig. 1C), the two upgrades (Fig. 1D) and the four-arm evaluation (Fig. 1E). We report the results in the same order: correspondence accuracy, identification of poorly matched cells, and downstream effects on data integration and label transfer.

### Optimal-transport framework for cross-modal correspondence

We first measured how accurately each method recovered true cell pairs. This was possible in co-assay data because both modalities were measured in the same cells. We evaluated each coupling using the fraction of samples closer than the true match ((12)), averaged across the 29 co-assay pairs. A value of 0 means every cell is closest to its true partner, whereas 0.5 indicates random matching. FOSCTTM cannot be applied to diagonal pairs because no true pairing exists. We compared eight couplings: four developed here (Diffusion-OT, CCA-OT, Diffusion-GW and Ensemble-OT) and four published methods (MaxFuse, SCOT, UnionCom and MMD-MA). All were tuned on designated co-assay datasets for each modality pair. The known pairing was used only to evaluate candidate settings and not by the algorithms themselves (Methods).

The couplings use different sources of information. Diffusion-OT compares cells through neighbourhood structure using diffusion coordinates from each modality’s nearest-neighbour graph, with shared-feature profiles weighted most strongly to retain direct cross-modal evidence. CCA-OT combines shared-feature similarity with a Seurat-style anchor step (17) using mutual nearest-neighbour anchors to fit canonical correlation analysis. Diffusion-GW uses geometry alone by comparing within-modality distance patterns around each cell. Each cost matrix is converted into a balanced coupling using Sinkhorn optimal transport (Methods).

Diffusion-GW was substantially less accurate than the other methods (0.387; Supplementary Fig. S1), suggesting that geometry alone was insufficient for these datasets. We therefore retained it only as a geometry-based comparison. Diffusion-OT and CCA-OT had similar overall FOSCTTM values, 0.137 and 0.127 respectively (Fig. 2B), but differed by modality. Diffusion-OT performed better on RNA–methylome (0.253 vs 0.274), whereas CCA-OT performed better on RNA–ADT (0.075 vs 0.089) and RNA–ATAC (0.169 vs 0.176). We therefore retained both. MaxFuse performed best among the published methods, with an overall FOSCTTM of 0.137, compared with 0.281 for SCOT, 0.390 for MMD-MA and 0.485 for UnionCom. It ranked second to Ensemble-OT on RNA–ADT (0.061) but performed weakest on RNA–methylome (0.370). Its joint embedding was converted to a distance matrix and then to a balanced coupling (Methods).

**Figure 2.**
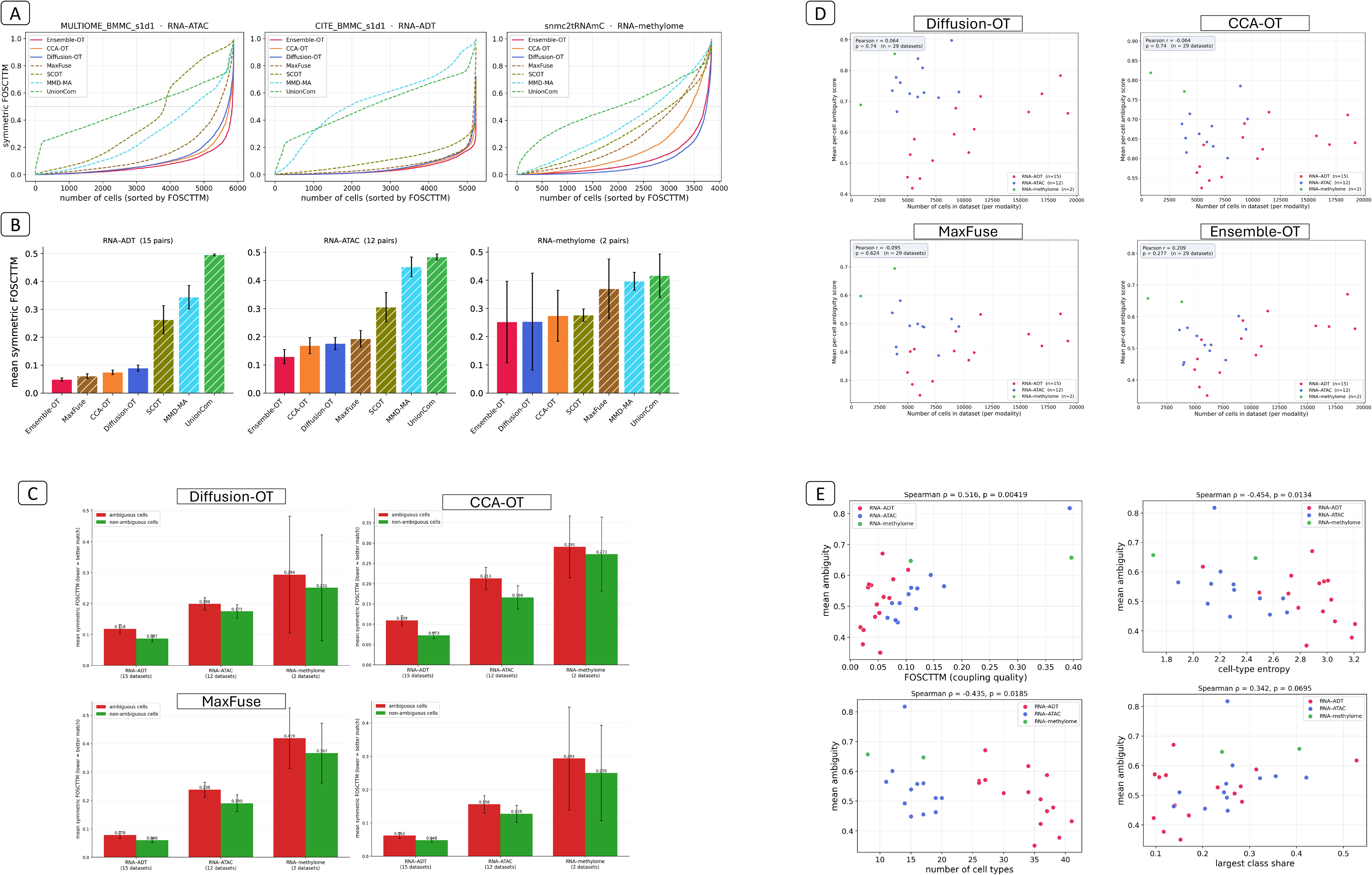
Cross-modal correspondence quality, ambiguity, and dataset characteristics in co-assay multi-omics data. (A) Per-cell symmetric FOSCTTM curves for one representative dataset of each co-assay modality pair: MULTIOME_BMMC_s1d1 for RNA–ATAC, CITE_BMMC_s1d1 for RNA–ADT and snmc2tRNAmC for RNA–methylome. Cells are ordered by FOSCTTM along the horizontal axis, and the seven curves are Ensemble-OT, CCA-OT, Diffusion-OT, MaxFuse, SCOT, MMD-MA and UnionCom. Lower values are better. (B) Mean symmetric FOSCTTM across all co-assay dataset pairs, in one sub-panel per modality pair: RNA–ADT (15 pairs), RNA–ATAC (12) and RNA–methylome (2). Within each sub-panel the seven couplings are ordered from most to least accurate, so the order differs between sub-panels. Error bars show one standard deviation across datasets. (C) Mean FOSCTTM for cells classified as ambiguous (red) and non-ambiguous (green). Each sub-panel is one coupling, and within it the three groups are the three co-assay modality pairs. Error bars show one standard deviation across datasets. (D) Dataset size against mean ambiguity, one sub-panel per coupling. Each point is one co-assay dataset pair, coloured by modality pair, and each sub-panel reports the Pearson correlation with its p-value. (E) Mean ambiguity under Ensemble-OT against four dataset properties: coupling quality measured by FOSCTTM, cell-type entropy, number of annotated cell types, and the share of cells in the largest cell type. Each point is one co-assay dataset pair, coloured by modality pair, and each sub-panel reports the Spearman correlation with its p-value.

Because Diffusion-OT, CCA-OT and MaxFuse capture complementary information, we combined them into Ensemble-OT. Each method ranks candidate partners for every cell, and these rankings are combined using dataset-specific weights selected without known pairing or cell-type labels. The resulting cost matrix is then balanced using Sinkhorn optimal transport (Methods). For RNA–methylome, where MaxFuse was unavailable, Ensemble-OT combined only Diffusion-OT and CCA-OT. Ensemble-OT achieved the best overall FOSCTTM (0.096) and the best result in each modality pair: 0.049 for RNA–ADT, 0.129 for RNA–ATAC and 0.252 for RNA–methylome (Fig. 2B). We therefore retained Ensemble-OT, CCA-OT, Diffusion-OT and MaxFuse for subsequent analyses.

All methods performed worse on RNA–methylome than on RNA–ADT or RNA–ATAC (Fig. 2B), consistent with the more indirect relationship between methylation and gene expression (1, 31, 32). The sorted per-cell FOSCTTM curves showed that this was not driven by a small subset of poorly matched cells (Fig. 2A; Supplementary Fig. S1). Ensemble-OT remained low across most cells, whereas weaker methods increased progressively across the population. The improvement therefore reflected better matching across many cells rather than correction of only a small subset.

### Ambiguity scores capture unreliable cross-modal matches

We next quantified cell-level uncertainty in the inferred correspondence using an ambiguity score derived from the coupling. For each cell, cross-modal coupling mass was pooled by cluster or cell type and converted into group-level support. A dominant group yields low ambiguity, whereas similar support for the two leading groups yields high ambiguity. Scores were computed independently for cells in both modalities. RefA-Amb and RefB-Amb scored query cells against the labelled cell types of modality 1 or modality 2 and were used for label transfer. LF-Amb instead used clustering-derived pseudo-labels on both sides and was used for data integration, where labels were not used for scoring. The score was applied to all 78 dataset pairs.

Thresholds were selected from the elbow of each ranked ambiguity curve using the L-method (33). Before elbow detection, scores were transformed as exp(10a), where *a* is the ambiguity score, preserving the ranking while sharpening changes in curve shape. A 50% safeguard prevented more than half of the cells in either modality from being removed. This safeguard was triggered in only 10.4% of runs after transformation, leaving 89.6% of thresholds data-driven. An independent chord-distance rule agreed within two percentage points in 77.1% of transformed curves, compared with 35–43% before transformation (Supplementary Fig. 2). In the 29 co-assay datasets, where the true correspondence is known, ambiguous cells had consistently higher FOSCTTM than non-ambiguous cells across all four couplings. The same pattern held for the 5% most ambiguous cells in nearly every dataset and coupling (Fig. 2C), supporting the score as a marker of poor correspondence.

### Dataset properties and ambiguity score

Each dataset pair contained about 1,000–19,000 cells per modality, with a median of approximately 4,000. We analysed single-batch pairs to reduce confounding by technical batch variation and processed all 78 pairs from raw counts using the same pipeline to minimise preprocessing-related differences.

Mean ambiguity was largely independent of dataset size. Across co-assay pairs, the correlation between cell number and mean ambiguity ranged from −0.15 to +0.23 across couplings, and none of these correlations was statistically significant. Across diagonal pairs, correlations ranged from −0.33 to −0.06, with only one of eight coupling-by-regime tests reaching significance. Dataset size was therefore the weakest predictor of ambiguity (Fig. 2D).

Other dataset properties showed stronger associations. Across the 29 co-assay pairs with Ensemble-OT (Fig. 2E), higher FOSCTTM, indicating poorer recovery of the true correspondence, was associated with higher mean ambiguity (ρ = 0.516, p = 0.0042). Ambiguity decreased with increasing cell-type entropy (ρ = −0.454, p = 0.0134) and number of annotated cell types (ρ = −0.435, p = 0.0185). The proportion of cells in the largest type showed a weaker positive trend (ρ = 0.342, p = 0.0695), whereas the size of the smallest type was unrelated (ρ = 0.170, p = 0.379; not shown). In the 49 diagonal pairs, dataset size, number of cell types and smallest-class size were again unrelated to ambiguity. In contrast, ambiguity decreased with entropy (ρ = −0.301, p = 0.0354) and increased strongly with the proportion of cells in the largest class (ρ = 0.579, p = 1.31 × 10⁻⁵; Supplementary Fig. 3). Overall, ambiguity was associated more strongly with correspondence quality and cell-type composition than with dataset size, with cell-type imbalance showing the clearest effect in diagonal data.

### Coupling upgrade and ambiguity-guided removal refine data integration

NMI, ARI and cell-type silhouette measure preservation of biological structure, whereas batch ASW, kBET and integration LISI measure cross-modal mixing (7, 20, 34). These objectives do not always improve together. Complete mixing can destroy biological structure, whereas complete separation preserves structure but fails to integrate the modalities (7). Among the mixing metrics, batch ASW and kBET provide more independent evidence because LISI is optimized directly by the upgrade. Each tool and coupling was tested separately by comparing upgraded and baseline values across datasets within each modality pair using Wilcoxon signed-rank tests. This yielded 40 tests per diagonal group and 44 per co-assay group, except for co-assay mixing metrics, where WNN was not applicable.

The upgrade consistently improved mixing and improved biological structure in several settings. iLISI increased significantly in 52.5–92.5% of tests across modality pairs, and kBET increased significantly in 32.5–90.0%. Neither metric showed a significant decrease. Batch ASW, the most conservative metric (7), increased significantly in 35.0–42.5% of tests across the three diagonal groups and in 40.0% of tests for co-assay RNA–ATAC. Biological structure improved in both RNA–ATAC groups and in co-assay RNA–ADT. NMI increased significantly in 45.0–52.3% of tests and cell-type silhouette in 45.5–62.5%. In contrast, diagonal RNA–methylome and diagonal RNA–RNA showed improved mixing without corresponding gains in structure. Co-assay RNA–ADT showed a trade-off: biological structure improved, whereas batch ASW decreased significantly in 50.0% of tests. Co-assay RNA–methylome contained only two datasets and was not suitable for significance testing. These summaries include all datasets. Safe-upgrade accepted no correction in 28.6% of all tool-by-coupling-by-dataset runs and returned the original embedding instead. This occurred in 29.6% of diagonal runs and 27.1% of co-assay runs.

Removing ambiguous cells improved biological structure more than removing the same number of cells at random. Relative to the matched random-removal control, NMI increased by 0.0318, ARI by 0.0368 and cell-type silhouette by 0.0059. These metrics improved in 39 of 40 tool-by-coupling tests and were significantly better in 95–98% of tests, with no significant decreases. The gains were also larger than those produced by the coupling upgrade alone. Relative to the original embedding, ambiguity-guided removal increased NMI by 0.0429 and ARI by 0.0477, compared with increases of 0.0121 and 0.0134 after the upgrade alone. Random removal increased NMI by only 0.0111 and ARI by 0.0109. Mixing changed little after ambiguity-guided removal, with batch ASW decreasing by 0.0040 and LISI changing by −0.0004. The largest structural gains were observed in diagonal RNA–ATAC, where NMI increased by 0.0388 and ARI by 0.0485, and cell-type silhouette improved in all 40 tests. Mixing in this group changed little. Significance was assessed using Wilcoxon signed-rank tests on per-dataset differences for each method, coupling and modality group. Removal was applied to the embedding returned by Safe-upgrade. When no correction was accepted, cells were removed from the original embedding.

The effect of ambiguity-guided removal differed between diagonal and co-assay data. In the diagonal groups, biological structure improved with little change in modality mixing. Diagonal RNA–RNA showed the most consistent improvement. Cell-type silhouette increased by 0.0099 and was significantly higher in all 40 tests. This group was also the only one in which mixing improved, with LISI increasing by 0.0145. Diagonal RNA–methylome showed a similar pattern, but only six dataset pairs were available. With this sample size, the smallest possible Wilcoxon p-value is 0.031, which limited the number of significant results.

In the co-assay groups, improved biological structure was accompanied by reduced modality mixing. NMI increased by 0.0307 in RNA–ADT and was significantly higher in all 44 tests. It also increased by 0.0348 in RNA–ATAC. However, batch ASW and LISI decreased in both groups, with larger reductions in RNA–ATAC of 0.0062 and 0.0143, respectively. Co-assay RNA– methylome contained only two datasets and was therefore not suitable for significance testing. At the individual-dataset level, ambiguity-guided removal outperformed random removal in 57– 67% of comparisons for biological structure, but only 31–43% for modality mixing (Supplementary Fig. 4). Full group-level results are provided in Data_Integration_supplementary.xlsx.

This difference may arise because ambiguous cells often occupy intermediate positions between both cell types and modalities. Removing these cells makes cell-type groups more compact and therefore improves NMI, ARI and silhouette, but it can also reduce overlap between modalities. This effect was stronger in co-assay data, where the two modalities were already paired at the cell level. In diagonal data, the inferred coupling may partly preserve cross-modal alignment after ambiguous cells are removed.

### Integration upgrade and ambiguity-guided removal depend on the tool and coupling choice

The previous analyses pooled tools and couplings within each modality group. This showed the overall effect of ambiguity-guided removal, but not which tools benefited most or which coupling performed best. We therefore examined the upgrade and removal separately for each tool and coupling. Figures 3A and 3B show the ten tools applicable to diagonal RNA–ATAC, including the three encoder-based models developed here. Results for the other groups are shown in Supplementary Fig. 4. Each bar is the mean across all datasets in the group, including those for which Safe-upgrade returned the original embedding. Ambiguous cells were removed from the embedding returned by Safe-upgrade, whether corrected or original. WNN was included only for co-assay data because it requires both modalities to be measured in the same cells. Individual results are highlighted for the two best published tools and the best model developed here, ranked by final performance.

**Figure 3.**
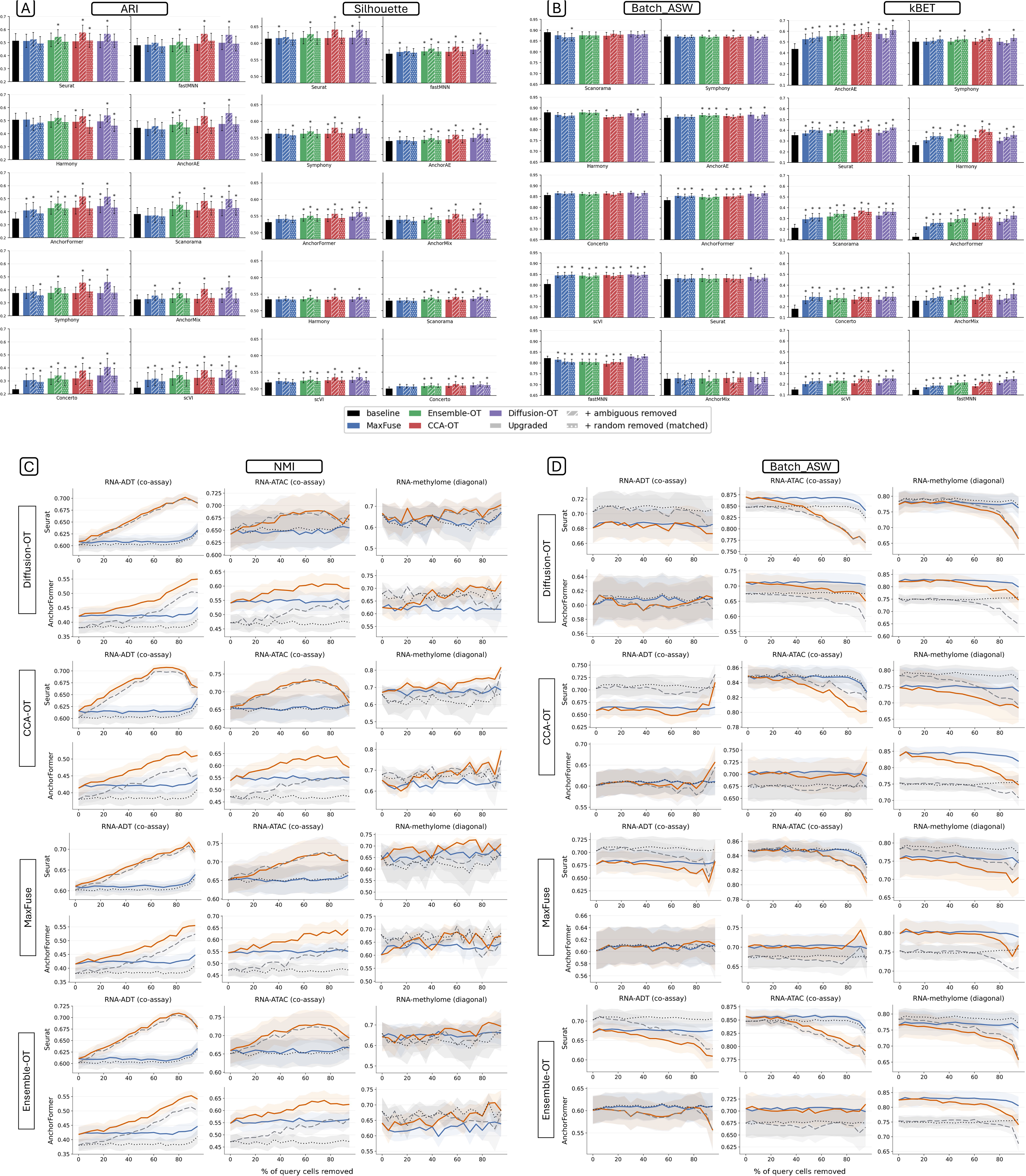
Effect of the coupling-based embedding upgrade and of ambiguity-guided cell removal on integration quality. (A, B) Bar charts for the diagonal RNA-ATAC group (25 dataset pairs), scored by cell-type structure in A (ARI and cell-type silhouette) and by modality mixing in B (batch ASW and kBET). Each panel is one of ten integration tools and holds thirteen bars: the original embedding (black), then MaxFuse, Ensemble-OT, CCA-OT and Diffusion-OT, each upgraded (solid), upgraded with ambiguous cells removed (diagonal hatching), and upgraded with the same number of random cells removed (dotted hatching). WNN is not shown, as it requires both modalities in the same cells. Bars give the mean across datasets and error bars one standard error; an asterisk marks p < 0.05 against that tool’s own baseline (Wilcoxon signed-rank test on paired per-dataset differences). (C, D) The same comparison across the removal sweep, scored by NMI in C and batch ASW in D. Each block is one coupling, in the order Diffusion-OT, CCA-OT, MaxFuse and Ensemble-OT; within a block the rows are Seurat and AnchorFormer and the columns are co-assay RNA-ADT (15 pairs), co-assay RNA-ATAC (12) and diagonal RNA-methylome (6). The horizontal axis is the percentage of cells removed, from 0 to 95 in steps of 5. Four conditions are drawn: ambiguity-guided removal on the upgraded embedding (orange, solid) and on the original embedding (grey, dashed), and matched random removal on the upgraded embedding (blue, solid) and on the original embedding (black, dotted). Shaded bands give one standard error across datasets.

Among the five groups with sufficient datasets, the contributions of the upgrade and removal differed. In diagonal RNA–ATAC, the upgrade improved mixing for every tool, but by different amounts. iLISI increased from 0.0885 to 0.1393 for AnchorAE but only from 0.0137 to 0.0182 for Seurat. Ambiguity-guided removal increased NMI by 0.0334–0.0399 over matched random removal and outperformed the control in 81–89% of comparisons. Coupling choice affected removal more than the upgrade, with an NMI spread of 0.070 for removal compared with 0.031 for the upgrade. A similar pattern appeared in the other groups. Diagonal RNA–methylome was the only group in which improved mixing was accompanied by lower structure. Concerto gained 0.2076 in batch ASW, while NMI decreased slightly under every coupling. Only six datasets were available, which limited statistical significance. In diagonal RNA–RNA, baseline NMI was already high at 0.8313–0.8784. The upgrade slightly reduced structure, whereas removal still improved NMI by 0.0122–0.0285 over the matched control. This was also the only group in which removal improved mixing.

Co-assay RNA–ADT showed the most consistent removal benefit, outperforming matched random removal in 90–100% of comparisons. Co-assay RNA–ATAC showed the largest mixing gains from the upgrade, with LISI increasing by 0.1205–0.1293. Random removal also increased NMI in this group, including a gain of 0.0215 for fastMNN, showing why the matched control was necessary. WNN differed from the other tools in both co-assay groups. Batch ASW decreased by 0.0245 in RNA–ADT and 0.0364 in RNA–ATAC.

Across all dataset pairs, the upgrade improved mixing for every applicable tool, although the gains varied substantially. LISI increased by 0.1601 for AnchorFormer and 0.1490 for Scanorama, compared with 0.0313 for Seurat and 0.0033 for Symphony. Changes in structure were smaller and inconsistent. NMI increased by 0.0474 for scVI but decreased by 0.0097 for Harmony. WNN remained unchanged because Safe-upgrade requires an increase in LISI, which was already at its maximum. Tools with weaker baseline embeddings therefore tended to gain more from the upgrade. Coupling choice had relatively little effect, with NMI differences below 0.02 in 65% of tool-by-group combinations.

For ambiguity-guided removal, coupling choice mattered more. Relative to matched random removal, CCA-OT increased NMI by 0.0462, whereas MaxFuse increased it by 0.0143. CCA-OT performed best in five of the six modality groups, while MaxFuse performed worst in all six. Differences across integration tools were smaller, with NMI gains ranging from 0.0215 to 0.0413. Only Harmony with MaxFuse failed to outperform the matched control, with a difference of −0.0001.

The two uses of the coupling therefore had different roles. The upgrade mainly improved modality mixing and was influenced more by the integration tool than by the coupling. Ambiguity-guided removal mainly improved biological structure and was influenced more strongly by the coupling used to identify ambiguous cells.

### Ambiguity-guided removal improves cell-type structure with limited loss of mixing

Figures 3C and 3D extend the analysis across removal levels from 5% to 95%. The main figure shows Seurat and AnchorFormer for NMI (Fig. 3C) and batch ASW (Fig. 3D) in co-assay RNA– ADT, co-assay RNA–ATAC and diagonal RNA–methylome. Results for the other groups and tools are shown in Supplementary Fig. 5. At 30% removal, the upgrade remained beneficial in both regimes, with the largest gain in co-assay RNA–ATAC and the smallest in diagonal RNA– RNA, where baseline performance was already high. Ambiguity-guided removal outperformed matched random removal in all 24 group-by-coupling comparisons. The additional NMI gain ranged from 0.0006 to 0.0656. Diagonal RNA–methylome showed the smallest gains, with three couplings adding no more than 0.0029, whereas CCA-OT added 0.0340.

Across the removal sweep, random removal had little effect on NMI. Between 5% and 75% removal, its mean change ranged from 0.0035 to 0.0134. In contrast, the advantage of ambiguity-guided removal increased progressively. The mean NMI difference from the matched control increased from 0.0050 at 5% removal to 0.0380 at 30% and 0.0619 at 50% (Fig. 3C). This pattern indicates that the gain depended on which cells were removed rather than removal alone. Seurat and AnchorFormer showed similar gains at 30% removal, with NMI increases of 0.0294 and 0.0279, respectively.

The improvement in cell-type structure was accompanied by a gradual reduction in modality mixing. Relative to matched random removal, batch ASW decreased by 0.0038 at 30% removal, 0.0131 at 50% and 0.0196 at 80% (Fig. 3D). At 50% removal, diagonal RNA–RNA showed both the largest NMI gain (0.0716) and the largest batch ASW decrease (0.0507). At lower removal levels, however, structural gains appeared before substantial changes in mixing. At 20% removal in co-assay RNA–ATAC, NMI increased by 0.0253 while batch ASW changed by only 0.0029. The elbow rule used in Figures 3A and 3B removed a median of 30.3% of cells, ranging from 17.6% to 41.1% across groups.

When the upgrade and removal were considered together, CCA-OT performed best in five of the six modality groups. Relative to the original embedding with matched random removal at 30%, CCA-OT increased NMI by 0.0867 in diagonal RNA–ATAC, 0.0819 in co-assay RNA– ATAC, 0.0719 in diagonal RNA–RNA, 0.0587 in co-assay RNA–ADT and 0.0384 in diagonal RNA–methylome. Co-assay RNA–methylome was the exception, where Diffusion-OT achieved the largest gain (0.0502 versus 0.0348 for CCA-OT). MaxFuse performed worst in both diagonal RNA groups and produced no gain in co-assay RNA–methylome. CCA-OT also outperformed the matched control in 96.1% of comparisons for diagonal RNA–RNA, 97.6% for co-assay RNA–ADT, 90.8% for diagonal RNA–ATAC and 68.3% for diagonal RNA–methylome. Ensemble-OT performed slightly better in the two remaining co-assay groups. Diagonal RNA– methylome was the least consistent setting, where Diffusion-OT and MaxFuse outperformed the control in only about half of comparisons.

These results support CCA-OT as a practical default when the upgrade and ambiguity-guided removal are used together. The upgrade improves cross-modal mixing, while removal mainly improves cell-type structure.

### UMAP visualizations highlight ambiguous populations and changes in embedding structure

Figure 4 visualizes the four conditions quantified in Figure 3. Each row shows one integration tool applied to one dataset, including the original embedding, the upgraded embedding, and the upgraded embedding after removing either ambiguous cells or the same number of randomly selected cells. Cells are coloured by cell type, and the table below each panel reports the size of each population and the proportion flagged as ambiguous. Four datasets are shown, with two from each regime: MULTIOME_CORTEX, human fetal cerebral cortex (35) (co-assay RNA– ATAC, MaxFuse, 26.6% flagged); CITE_BMMC_s1d3, human bone marrow mononuclear cells (3) (co-assay RNA–ADT, CCA-OT, 19.2%); shareP7RNAATAC, mouse skin and hair follicle at postnatal day 7 (36) (diagonal RNA–ATAC, CCA-OT, 47.0%); and mopF004×3CRNAmC, adult mouse primary motor cortex (31, 37) (diagonal RNA–methylome, Ensemble-OT, 33.4%).

**Figure 4.**
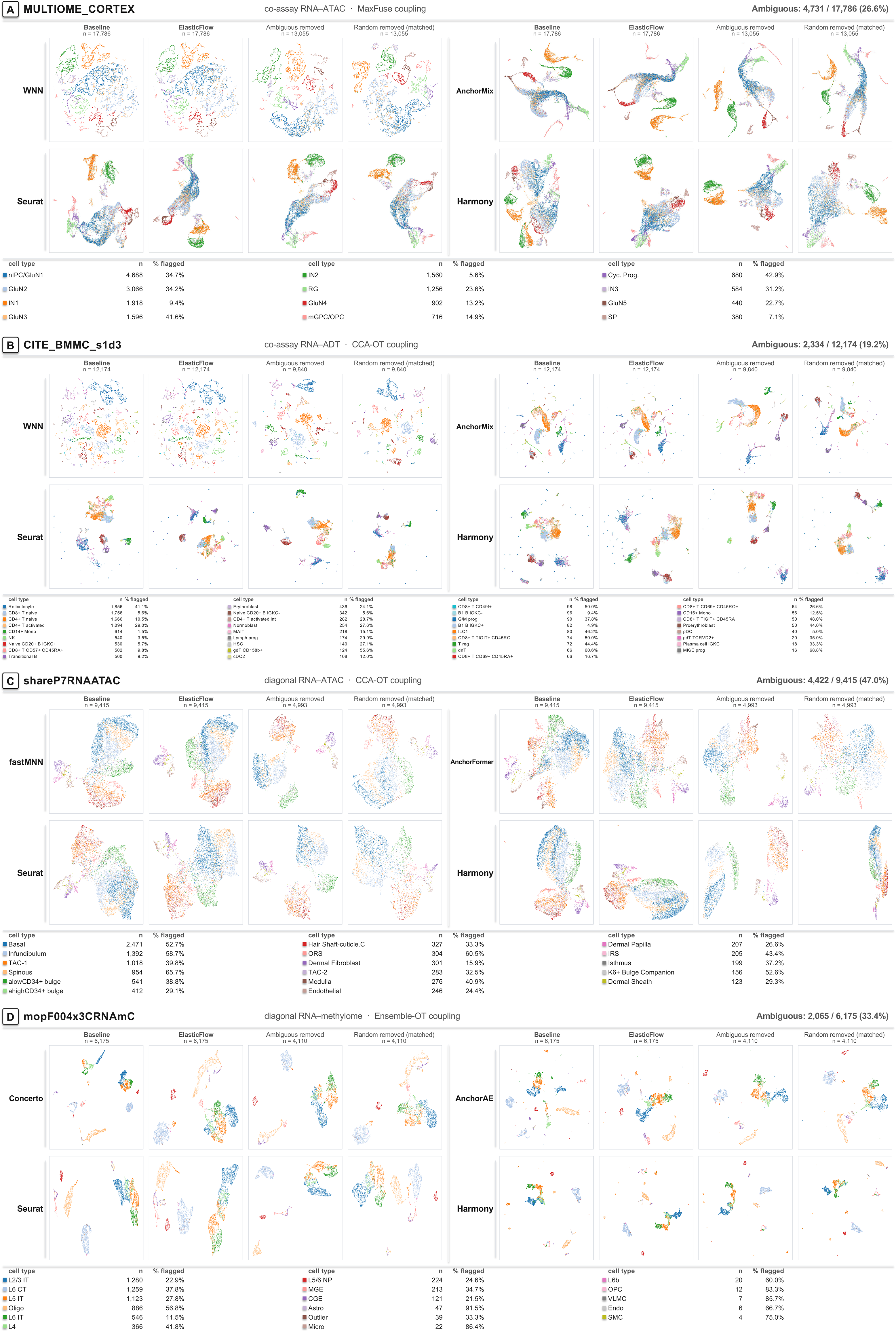
UMAP views of the embedding before and after the coupling-based upgrade and after ambiguity-guided cell removal. Four datasets are shown, two from each regime: (A) MULTIOME_CORTEX, a co-assay RNA-ATAC pair coupled with MaxFuse, 4,731 of 17,786 cells flagged as ambiguous (26.6 percent); (B) CITE_BMMC_s1d3, co-assay RNA-ADT with CCA-OT, 2,334 of 12,174 (19.2 percent); (C) shareP7RNAATAC, diagonal RNA-ATAC with CCA-OT, 4,422 of 9,415 (47.0 percent); and (D) mopF004×3CRNAmC, diagonal RNA-methylome with Ensemble-OT, 2,065 of 6,175 (33.4 percent). Each panel shows four integration tools in a two-by-two block: WNN, AnchorMix, Seurat and Harmony in A and B; fastMNN, AnchorFormer, Seurat and Harmony in C; and Concerto, AnchorAE, Seurat and Harmony in D. WNN appears only in the co-assay panels, as it requires both modalities in the same cells. Each tool is drawn as a row of four UMAPs in the same order: the original embedding, the embedding after the ElasticFlow upgrade, that upgraded embedding after the cells flagged as ambiguous are removed, and the upgraded embedding after the same number of cells is removed at random, matched per modality. Only the third and fourth images in a row hold the same number of cells. The number of cells retained is given above each UMAP. Points are single cells coloured by annotated cell type, and the colour key below each panel lists every cell type, the number of cells it contains and the percentage of it that was flagged.

The relative effects of the upgrade and removal differed across datasets. In diagonal RNA– ATAC, ambiguity-guided removal produced the larger change. For AnchorFormer on shareP7RNAATAC, the upgrade increased NMI from 0.3785 to 0.4222, while removal increased it further to 0.5380. The other three tools showed similar gains over matched random removal. In diagonal RNA–methylome, the upgrade contributed more. For Concerto, batch ASW increased from 0.6255 to 0.8160 as the scattered embedding became more integrated, while removal added little further improvement. Both steps contributed to the co-assay examples, with a larger upgrade effect in MULTIOME_CORTEX and a larger removal effect in CITE_BMMC_s1d3. WNN was unchanged by the upgrade because Safe-upgrade could not increase an already maximal mixing score. After removal, however, batch ASW decreased by 0.0281 and 0.0614 in the two co-assay datasets.

The cell-type tables show that ambiguous cells were not distributed evenly across populations. In CITE_BMMC_s1d3, more than half of MK/E progenitors (68.8%, n = 16), dnT cells (60.6%, n = 66) and gdT CD158b+ cells (55.6%, n = 124) were flagged, compared with 1.5% of CD14+ monocytes and 3.5% of NK cells. In MULTIOME_CORTEX, 42.9% of cycling progenitors and 41.6% of GluN3 cells were flagged, compared with 5.6% of IN2 cells. In shareP7RNAATAC, the corresponding proportions were 65.7% for spinous cells, 60.5% for outer root sheath cells and 15.9% for dermal fibroblasts. In mopF004×3CRNAmC, flagging was especially high among several rare populations, including astrocytes (91.5%, n = 47), microglia (86.4%, n = 22), VLMC cells (85.7%, n = 7) and OPCs (83.3%, n = 12), compared with 11.5% of L6 IT cells (n = 546). Oligodendrocytes were an exception, with 56.8% flagged despite being common. These examples show that ambiguity-guided removal preferentially affects populations that are less clearly resolved across modalities rather than removing a fixed proportion from each cell type.

UMAP provides only a two-dimensional view of the embeddings, so Figure 4 is intended to illustrate rather than quantify these effects (38). The visual patterns agree with the results in Figure 3. Ambiguous cells are concentrated in less well-resolved populations, and their removal generally produces more compact and separated cell-type groups. Same setting UMAPs color-coded with batches are presented Supplementary Fig. 6.

### Coupling upgrade and ambiguity-guided removal improve label transfer

We next tested whether coupling information improves cross-modal label transfer and whether removing cells with uncertain correspondence provides additional benefit. Ten label-transfer tools were evaluated across all 78 dataset pairs in both transfer directions and with each of the four couplings, using the same four conditions as for integration (Fig. 5). In the main analysis, RNA was the labelled reference, and the second modality was the unlabelled query. The reverse direction was also tested for every dataset. Performance was measured by accuracy and macro-F1, which gives equal weight to each cell type and is therefore more sensitive to rare populations. Query-cell ambiguity was calculated using only reference annotations, so query labels were never used to select cells for removal.

**Figure 5.**
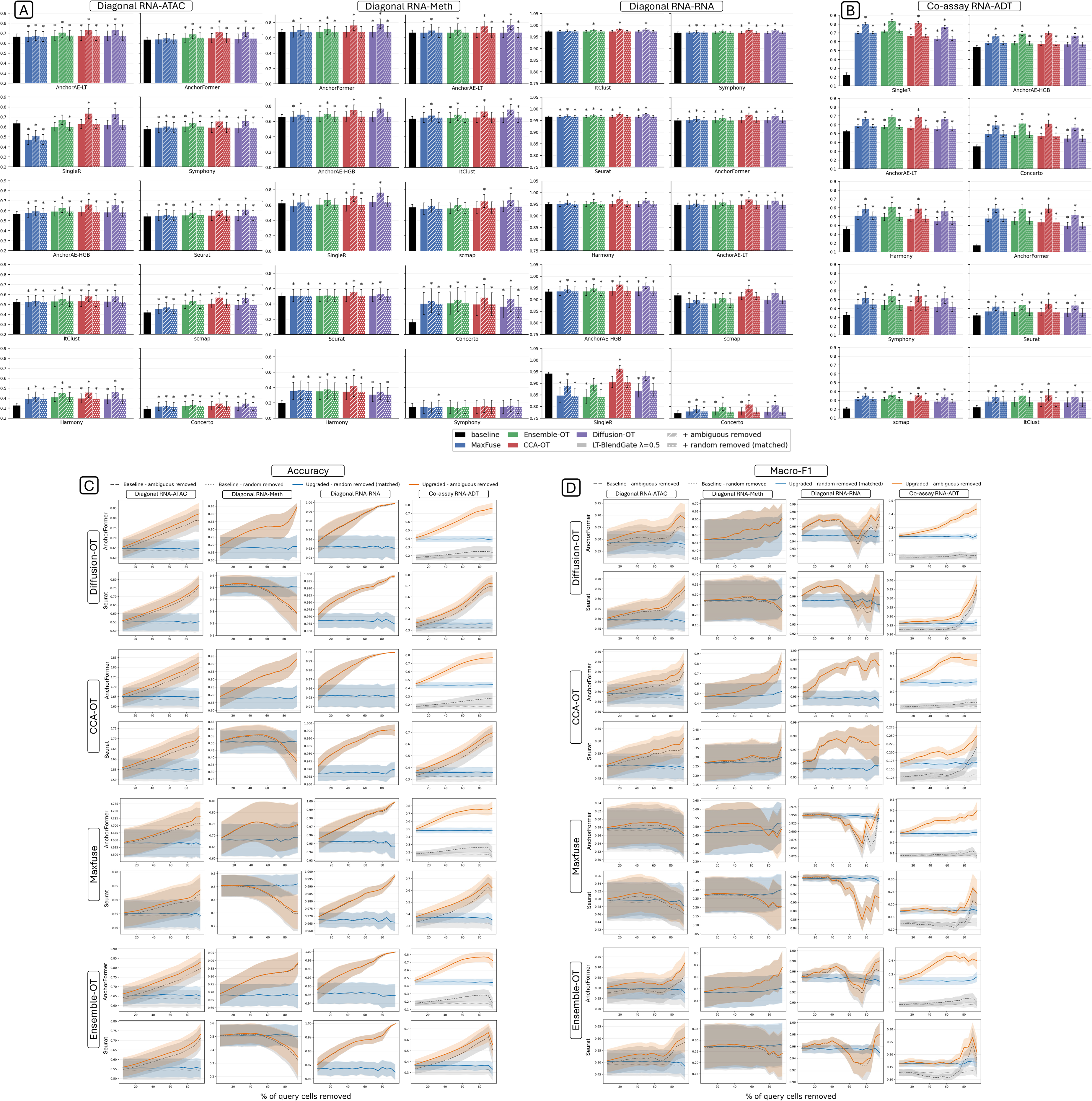
Effect of the coupling-based label-transfer upgrade and of ambiguity-guided removal of query cells. Labels are transferred from RNA to the second modality of each pair, and the upgrade is LT-BlendGate with λ = 0.5. (A) Mean accuracy for the three diagonal modality pairs: RNA-ATAC (25 dataset pairs), RNA-methylome (6) and RNA-RNA (18). (B) The same for the co-assay RNA-ADT pair (15). In both panels each small plot is one of ten label-transfer tools, ordered by decreasing baseline accuracy, and holds thirteen bars: the baseline (black), then MaxFuse, Ensemble-OT, CCA-OT and Diffusion-OT, each upgraded (solid), upgraded with ambiguous query cells removed (diagonal hatching), and upgraded with the same number of random query cells removed (dotted hatching). Bars give the mean across dataset pairs and error bars one standard error; an asterisk marks p < 0.05 against that tool’s own baseline (Wilcoxon signed-rank test on paired per-dataset differences). (C, D) The same comparison across the removal sweep, scored by accuracy in C and macro-F1 in D. Each has four blocks, one per coupling, in the order Diffusion-OT, CCA-OT, MaxFuse and Ensemble-OT; within a block the rows are AnchorFormer and Seurat and the columns are the four modality pairs of A and B, in that order. The horizontal axis is the percentage of query cells removed, from 5 to 95 in steps of 5, and each panel keeps its own vertical scale. Four curves are drawn: ambiguity-guided removal (orange) and matched random removal (blue) on the upgraded predictions, and the same two on the baseline predictions (grey dashed and grey dotted). Shaded bands give one standard error across dataset pairs.

Both the upgrade and ambiguity-guided removal improved performance. With RNA as the reference, the upgrade increased accuracy by 0.0412 and macro-F1 by 0.0343 across tools, datasets and couplings. After ambiguity-guided removal, the total gains reached 0.0948 and 0.0603. Among the 60 modality-group-by-tool tests, the upgrade significantly improved accuracy in 75% and macro-F1 in 67%. The combined treatment was significant in 93% and 85%, respectively. Random removal added almost no improvement beyond the upgrade, with an accuracy gain of 0.0410 above baseline compared with 0.0412 for the upgrade alone. Relative to this matched control, ambiguity-guided removal added 0.0538 accuracy and 0.0261 macro-F1 and outperformed random removal in 92.6% and 85.2% of paired comparisons.

The largest gains occurred in settings with lower baseline performance. Co-assay RNA–ADT had the lowest baseline accuracy (0.3279) and showed the largest improvement. The upgrade added 0.1338, and ambiguity-guided removal added a further 0.0986 over the matched control, outperforming it in 97.3% of comparisons. Across the four intermediate groups, with baseline accuracies of 0.4137–0.5206, the upgrade added 0.0172–0.0527 and removal added 0.0446– 0.0712 (Fig. 5A, B). Diagonal RNA–RNA had the highest baseline accuracy (0.9328). The upgrade reduced accuracy by 0.0090, but removal still added 0.0183 over the control and outperformed it in 94.2% of comparisons.

The effect also varied by label-transfer tool. SingleR showed the largest overall gain (0.1470) and the largest benefit from removal. Harmony benefited most from the upgrade, gaining 0.0844 in accuracy and 0.0945 in macro-F1. The models developed here showed smaller upgrade gains. AnchorAE and AnchorAE-HGB improved by only 0.0108 and 0.0134, respectively, but ambiguity-guided removal still added 0.0516 and 0.0550. Seurat showed the smallest gain from removal (0.0373).

Coupling performance also differed between the upgrade and removal. For the upgrade, MaxFuse performed best in co-assay RNA–ADT, increasing accuracy by 0.1512 compared with 0.1116 for Diffusion-OT. Ensemble-OT performed best in the other two co-assay groups and in diagonal RNA–methylome, while CCA-OT performed best in diagonal RNA–ATAC. None of the four couplings improved diagonal RNA–RNA, where accuracy decreased by 0.0031–0.0126 from an already high baseline. Differences among couplings were significant in five of the six modality groups by Friedman test (p < 0.001). Diagonal RNA–methylome was the exception, with all four couplings within 0.003 accuracy.

For ambiguity-guided removal, each coupling was compared with matched random removal to isolate the effect of cell selection. Diffusion-OT performed best in four groups, including both methylome groups, whereas CCA-OT performed best in the remaining two. All six comparisons were significant. MaxFuse performed worst in every group despite its strong upgrade performance in co-assay RNA–ADT. For the combined treatment, CCA-OT, Ensemble-OT and Diffusion-OT each performed best in two of the six groups. This differs from data integration, where CCA-OT was the strongest coupling across all six groups. Full results are provided in Label_transfer_supplementary.xlsx, with additional plots in Supplementary Fig. 7.

The reverse transfer direction produced similar results. Using the second modality as the reference, the upgrade increased accuracy by 0.0316 and macro-F1 by 0.0232. After ambiguity-guided removal, the total gains reached 0.0846 and 0.0382, compared with 0.0948 and 0.0603 when RNA was the reference. Relative to matched random removal, accuracy increased by 0.0531 and improved in 90.5% of comparisons, close to the 0.0538 gain and 92.6% win rate obtained with RNA as reference. The larger difference appeared in macro-F1. The gain decreased from 0.0261 to 0.0150, and the win rate from 85.2% to 74.3%, when the second modality was used as reference. This pattern suggests that rare cell types are more sensitive to the choice of reference modality. The weakest setting was diagonal RNA–methylome with methylome as the reference, where removal added 0.0345 and outperformed the control in 78.3% of comparisons. The strongest was co-assay RNA–ATAC with ATAC as the reference, with a gain of 0.0828 and a 97.9% win rate.

### Ambiguity-guided removal progressively improves label transfer

The benefit of ambiguity-guided removal increased as more cells were removed (Fig. 5C, D). Across removal levels from 5% to 95%, the matched random-removal curve remained nearly flat, with accuracy close to 0.610 up to 70% removal. In contrast, the advantage of ambiguity-guided removal increased from 0.0087 at 5% removal to 0.0539 at 30%, 0.0906 at 50% and 0.1479 at 80%. Macro-F1 showed the same pattern but with smaller gains, reaching 0.0261 at 30% removal and approximately 0.035–0.037 beyond 50%. Ambiguity-guided removal outperformed the matched control at every threshold for CCA-OT, Ensemble-OT and Diffusion-OT on both metrics. The only exception was MaxFuse, for which macro-F1 was 0.0037 lower than the control at 75% removal. The elbow threshold therefore balances performance gain against cell retention rather than identifying a minimum level at which removal becomes effective. Applied separately to each dataset and capped at 50%, it removed 27.0% of query cells on average. Removing more cells did not always produce a larger benefit. Diagonal RNA– RNA had the lowest removal rate but still outperformed random removal in 94.2% of comparisons, whereas co-assay RNA–methylome had the highest removal rate and outperformed it in 95.0%.

For accuracy, ambiguity-guided removal showed a progressively larger advantage in seven of the eight panels, while the random-removal curves remained nearly flat (Fig. 5C). Co-assay RNA–ADT showed the largest gain. At 30% removal, AnchorFormer exceeded random removal by 0.1343. Most of this gain came from the upgrade, with accuracy reaching 0.6177 after upgrading compared with 0.2053 without it. Diagonal RNA–RNA showed the smallest changes because baseline accuracy was already high, although the advantage of ambiguity-guided removal still increased across the sweep. The main exception was Seurat on diagonal RNA– methylome with MaxFuse. Its performance matched the control up to 30% removal but fell 0.0310 below it at 50%, consistent with the weaker cell-selection performance of MaxFuse.

Macro-F1 showed the same overall pattern, although gains were smaller and reached a plateau earlier (Fig. 5D). At 30% removal, ambiguity-guided removal outperformed the control in all eight panels. By 50%, the Seurat curves had largely plateaued, while accuracy continued to improve in the corresponding panels. Results for the remaining modality groups and tools are shown in Supplementary Fig. 8.

## Discussion

### Correspondence matrices capture alternative matches and coupling-specific ambiguity

A correspondence matrix is a soft transport plan in which each cell distributes its mass across possible partners in the other modality rather than selecting a single match (11, 12, 39). This is particularly important in diagonal data, where no unique physical partner exists. In co-assay data, the known cell pairing can be used to evaluate the coupling, whereas in diagonal data Γ represents plausible cross-modal relationships and ambiguity reflects competition among them (9).

The ambiguity score is derived from this distribution. A cell is considered ambiguous when its coupling mass is divided between competing cell-type groups. The score therefore measures uncertainty in correspondence rather than distance in a latent space. Because each coupling uses different information, they can identify different cells as ambiguous. CCA-OT relies on agreement between shared features, Diffusion-OT incorporates within-modality graph structure (40), MaxFuse uses graph smoothing and iterative pivot refinement for weakly linked assays (29), and Ensemble-OT combines all three. Moscot follows a related principle by using Wasserstein, Gromov–Wasserstein or fused Gromov–Wasserstein transport depending on the available cross-domain information (41).

The couplings also differed in the proportion of cells flagged as ambiguous. Across all 78 dataset pairs, Diffusion-OT flagged the largest proportion (36.4%), followed by CCA-OT (32.1%), Ensemble-OT (26.7%) and MaxFuse (24.8%). This ordering was observed in five of the six modality pairs. Co-assay RNA–methylome was the exception, where CCA-OT flagged 48.1% compared with 42.8% for Diffusion-OT. Ambiguity rates were highest in co-assay RNA– methylome (36–48%) and lowest in diagonal RNA–RNA (17–24%).

### Ambiguity is enriched in rare and transitional populations

Flagged cells were not distributed uniformly across cell populations. Rare, immature and transitional populations were frequently enriched for ambiguity, although the pattern differed between co-assay and diagonal data. In co-assay data, where the true cell pairing is known, ambiguity mainly reflects uncertainty between neighbouring cell states. In CITE_BMMC_s1d3, 68.8% of MK/E progenitors, 60.6% of dnT cells and 55.6% of gdT CD158b+ cells were flagged, compared with 1.5% of CD14+ monocytes and 3.5% of NK cells. This pattern is consistent with targeted antibody panels distinguishing mature lineages more clearly than closely related progenitor states (4).

Similar enrichment appeared in diagonal data, where no true cell-to-cell pairing exists. In developing cortex, 42.9% of cycling progenitors were flagged compared with 5.6% of IN2 cells. In skin, 65.7% of spinous cells were flagged compared with 15.9% of dermal fibroblasts. In the RNA–methylome example, 91.5% of astrocytes and 86.4% of microglia were flagged compared with 11.5% of L6 IT neurons. These examples span several tissues and suggest that ambiguity is not confined to a single biological system. A related pattern was reported by scOT-LT, where OT-based uncertainty highlighted query-specific populations absent from the reference (42). Ambiguity may therefore reflect genuine compositional mismatch as well as poor correspondence.

This interpretation is particularly relevant for immature and transitional populations. OT relates populations when direct pairings are unavailable, so cells occupying intermediate biological states may distribute transport mass across several neighbouring groups (9). High ambiguity does not necessarily indicate a low-quality cell or a failed alignment. It may instead identify regions where discrete labels or one-to-one correspondence provide an incomplete description of a biological continuum. In studies focused on transitional states, such cells may therefore be better retained for separate analysis than removed (9). Consistent with this interpretation, OT analysis of mouse pancreas development has recovered meaningful relationships among closely related developmental populations (41).

Dataset-level ambiguity was also related more strongly to correspondence quality and cell-type composition than to dataset size. Across the 29 co-assay pairs, correlations between cell number and mean ambiguity ranged from −0.15 to +0.23 across couplings, and none was statistically significant. In contrast, higher FOSCTTM, indicating poorer recovery of the true correspondence, was associated with higher mean ambiguity. Cell-type composition also mattered. Ambiguity decreased as cell-type entropy increased, while in diagonal data it increased when a single cell type accounted for a large proportion of cells. The number of annotated cell types and the size of the smallest population showed little association with ambiguity. These results suggest that ambiguity reflects correspondence quality and population imbalance more strongly than dataset size. In highly imbalanced datasets, the increased ambiguity is consistent with greater competition among similar candidate partners.

### Coupling upgrade and cell removal contribute differently to integration and label transfer

For data integration, the upgrade mainly improved modality mixing. The increase reached 0.1601 iLISI for AnchorFormer and 0.1490 for Scanorama, while changes in cell-type structure were smaller. Coupling choice had limited influence on this step, with differences below 0.02 NMI in 65% of tool-by-group combinations. Ambiguity-guided removal had a different effect. It improved cell-type structure by 0.0215–0.0413 NMI over matched random removal, with similar gains across tools but larger differences across couplings. This improvement was accompanied by reduced modality mixing. The decrease was small in the diagonal groups, where batch ASW changed by 0.0029–0.0095, and diagonal RNA–RNA was the exception, with iLISI increasing by 0.0145. The reduction in mixing was greater in co-assay data, where the modalities were already aligned at the cell level. In label transfer, both steps contributed substantially. With RNA as the reference, the upgrade increased accuracy by 0.0412 and ambiguity-guided removal added a further 0.0538 over matched random removal. Random removal added almost no improvement beyond the upgrade, with a total gain of 0.0410 compared with 0.0412 for the upgrade alone.

The coupling that performed best also depended on how it was used. For integration, coupling choice had little effect on the upgrade but strongly affected ambiguity-guided removal. CCA-OT performed best in five of the six modality pairs, whereas MaxFuse performed worst in all six. The difference was also clear in label transfer. MaxFuse produced the largest upgrade in co-assay RNA–ADT, increasing accuracy by 0.1512, but was the weakest coupling for cell selection in every modality group. Diffusion-OT showed the opposite pattern and performed best for cell selection in four of the six groups, including both methylome groups. No coupling was therefore optimal across all label-transfer settings, supporting selection according to both modality pair and downstream task (30, 43).

Transfer direction had little effect on accuracy but a larger effect on macro-F1. Relative to matched random removal, transfer from RNA increased accuracy by 0.0538 and macro-F1 by 0.0261. In the reverse direction, the gains were 0.0531 and 0.0150, while the macro-F1 win rate decreased from 85.2% to 74.3%. This suggests that rare populations are more sensitive to the choice of reference modality than common populations. The weakest setting was diagonal RNA–methylome with methylome as the reference, consistent with the lower resolution of closely related cell states in some non-transcriptomic assays.

The removal sweep showed a similar difference between accuracy and macro-F1. Accuracy continued to improve as more ambiguous cells were removed, reaching a gain of 0.1479 over matched random removal at 80%. Macro-F1 increased more modestly and plateaued at approximately 0.035–0.037 beyond 50% removal. Because ambiguous cells were enriched in rare populations, deeper removal may preferentially reduce representation of these classes. The benefit therefore increased gradually rather than appearing at a fixed threshold. The elbow criterion, which removed 27.0% of query cells on average, provides a compromise between improving prediction accuracy and retaining less abundant populations.

### Tool design shapes the benefit of coupling-based refinement

The benefit of coupling-based refinement depended on each tool’s design. Concerto learns a contrastive embedding without estimating a transport map (27), while Symphony maps query cells into a fixed reference (21); both showed relatively large gains from ambiguity-guided removal. Seurat filters and scores weak anchors before label transfer (17), and Harmony performs correction at the cluster level (20), leaving less room for additional refinement. Our models followed their intended settings. AnchorMix, designed for co-assay data, was the strongest custom model in that regime, reaching 0.6158 in RNA–ADT and 0.7001 in RNA– ATAC by using the known pairing directly. AnchorAE and AnchorFormer were designed for diagonal data and performed best there. AnchorAE achieved the highest final score in diagonal RNA–methylome (0.7252), while AnchorFormer showed the largest mixing gain overall (0.1601 iLISI). Both use reciprocal nearest-neighbour anchors during training (19), which may partly explain their smaller gains from additional coupling-based filtering. WNN showed the clearest dependence on known pairing. It constructs a joint representation from paired modalities (23), so Safe-upgrade could not improve its already maximal mixing score. Ambiguity-guided removal, however, produced the largest reduction in mixing for WNN because removing paired cells disrupted this structure.

### Limitations

Several limitations qualify our conclusions. First, integration LISI should be interpreted cautiously because Safe-upgrade directly optimizes this metric. Batch ASW and kBET therefore provide more independent evidence and generally showed smaller gains. Second, benefits depended on both the task and the metric. Improvements in cell-type structure sometimes reduced modality mixing, and the coupling best suited to embedding correction was not always best for ambiguity-guided removal. Third, Safe-upgrade returns the original embedding when no candidate satisfies its criteria. With Diffusion-OT, this occurred in 63.3% of Seurat and 20.4% of Harmony diagonal cases, compared with 6.9% and 3.4% in co-assay data. Ambiguity also depends on the coupling and pseudo-label detector, so different choices may identify different cells.

The random-removal control matched the number of removed cells but not their cell-type composition. Because ambiguous cells were enriched in rare and transitional populations, some gains may result from removing intrinsically difficult populations rather than improving the representation of the remaining cells. In studies focused on these populations, flagging them for separate analysis may therefore be more appropriate than removing them.

All dataset pairs also shared the same complete cell-type label set across modalities. We therefore did not test cases in which a population is present in one modality but absent from the other. In such settings, cells from the missing population would be forced to distribute coupling mass among the available reference types and may be flagged as ambiguous. Whether the score can distinguish this type of compositional mismatch from a genuinely transitional state remains unknown and would require experiments in which cell types are deliberately withheld.

OT has also been applied beyond cross-modal integration. Moscot scaled correspondence analysis to 1.7 million embryonic cells across 20 time points and incorporated temporal, multimodal and spatial information (41). Extending our framework to such settings could test whether ambiguity also identifies uncertain developmental transitions or regions of unreliable downstream inference. Neural OT may further allow transport maps to generalize to unseen cells and perturbation responses (44). Our framework currently measures correspondence only between observed modalities, so these temporal and perturbational settings remain to be tested (9). Recommended tool–coupling combinations from the settings evaluated here are summarized in Table 1. Each suggestion is the combination that scored highest after the upgrade and ambiguity-guided removal, by NMI for integration and by accuracy for label transfer, averaged over the datasets of each modality pair. If modality mixing matters more for a given application, other combinations would be chosen. The final choice is therefore a trade-off between conserving biological structure and mixing the two modalities.

**Table 1.** Suggested tool and coupling for each modality pair. (A) Data integration and (B) label transfer. Rows are the benchmarked tools and columns the four correspondence matrices. A star marks the tool-and-coupling combination we suggest for that modality pair, chosen by the final score after the full treatment, that is the coupling upgrade followed by ambiguity-guided removal. Where the two best combinations in a pair lie within 0.002 of each other, both carry a star, because a gap that small does not express a real preference. Only rows carrying a star are shown; the full tables, with the best coupling for every tool, are in the supplementary file. Integration is ranked by NMI and label transfer by accuracy with RNA as the reference. The same coupling also supplies the ambiguity score for that pair. These are suggestions drawn from the present benchmark rather than established defaults, and they are averages over the datasets of each group, so an individual dataset may depart from them. WNN appears only in the co-assay sections, because it requires both modalities to be measured in the same cells.

A. Data integration
| Tool | CCA-OT | Ensembl e-OT | Diffusion-OT | MaxFuse |
| --- | --- | --- | --- | --- |
| **Co-assay RNA-ADT** |  |  |  |  |
| WNN | ★ |  |  |  |
| AnchorMix |  | ★ |  |  |
| **Co-assay RNA-ATAC** |  |  |  |  |
| WNN | ★ |  |  |  |
| AnchorMix |  |  | ★ |  |
| **Co-assay RNA-methylome** |  |  |  |  |
| WNN |  |  | ★ |  |
| **Diagonal RNA-ATAC** |  |  |  |  |
| fastMNN | ★ |  |  |  |
| **Diagonal RNA-methylome** |  |  |  |  |
| AnchorAE |  |  | ★ |  |
| **Diagonal RNA-RNA** |  |  |  |  |
| Seurat | ★ |  | ★ |  |

B. Label transfer
| Tool | CCA-OT | Ensembl e-OT | Diffusion-OT | MaxFuse |
| --- | --- | --- | --- | --- |
| **Co-assay RNA-ADT** |  |  |  |  |
| SingleR |  | ★ |  |  |
| **Co-assay RNA-ATAC** |  |  |  |  |
| SingleR | ★ | ★ |  |  |
| **Co-assay RNA-methylome** |  |  |  |  |
| AnchorAE | ★ |  |  |  |
| **Diagonal RNA-ATAC** |  |  |  |  |
| SingleR | ★ |  |  |  |
| AnchorAE |  |  | ★ |  |
| **Diagonal RNA-methylome** |  |  |  |  |
| AnchorFormer |  |  | ★ |  |
| **Diagonal RNA-RNA** |  |  |  |  |
| ItClust | ★ |  |  |  |

### Conclusion

Making cross-modal correspondence explicit provides a per-cell measure of how reliably cells are matched across modalities. Across 78 dataset pairs and six modality pairings, evaluated with eleven integration and ten label-transfer tools, incorporating the coupling into existing methods substantially improved label transfer and improved data integration mainly through better modality mixing. The most consistent benefit came from ambiguity scoring. Ambiguous cells were more poorly matched in almost every co-assay dataset, and their removal outperformed matched random removal in 39 of 40 integration tests and 92.6% of label-transfer comparisons. These cells were enriched in rare, immature and transitional populations, indicating that ambiguity often concentrates where cross-modal correspondence is least clearly resolved. Coupling choice had little effect on embedding correction but strongly influenced ambiguity-guided cell selection, and no single coupling was optimal across all tasks and modality pairs. Overall, reporting the correspondence and its uncertainty provides information that is lost when the coupling remains hidden inside the analysis pipeline.

## Methods

This section presents the equations required to follow the main analyses. Full derivations, additional formulas, parameter grids and descriptions of all benchmarked tools are provided in the Supplementary Methods (Methods_supplementary.pdf).

### Preprocessing Datasets

We assembled 78 dataset pairs spanning six modality pairings from sixteen published studies (3, 35–37, 45–55). All pairs were processed from raw counts using the same pipeline to minimise preprocessing-related variation. RNA matrices were processed with Seurat SCTransform (56), retaining cells with 100–10,000 detected genes and fewer than 30% mitochondrial reads. RNA– ATAC data were represented in gene space by summing peaks overlapping each gene body and 2 kb upstream using Ensembl annotations (57), followed by log normalization. ADT counts were normalized using a centered log-ratio transformation across cells (4), and protein names were mapped to gene symbols. RNA–methylome pairs were converted to gene-level activity using ALLCools (31) and inverted so that larger values correspond to higher expected activity in both modalities. All normalized matrices were mapped to gene symbols, giving the two modalities a common feature vocabulary. Co-assay pairs retained their measured cell pairing, whereas no cell-level pairing was assumed for diagonal pairs. Full filtering criteria, thresholds, feature counts and source studies are provided in the Supplementary Methods.

### Correspondence Matrix Inference with Optimal Transport

We benchmarked four published correspondence methods, MaxFuse (29), SCOT (12), MMD-MA (16) and UnionCom (58), together with four methods developed here. Methods were tuned separately for each modality pair using designated co-assay datasets, where the measured pairing provided a direct tuning criterion. The selected configuration was then applied unchanged to the remaining datasets of the same modality pair. Search grids and selected settings are provided in the Supplementary Methods.

Each method produces a cross-modal cost matrix *C* ∈ ℝ*^n^*^1×*n*2^, where *n*_1_ and *n*_2_ are the numbers of cells in the two modalities. Smaller *C_i_*_j_ values indicate stronger candidate correspondence between cell *i* and cell *j*. Except where otherwise specified, the resulting cost is converted into a soft balanced coupling using the entropic transport procedure described below.

Diffusion-OT combines within-modality manifold structure with direct cross-modal feature similarity (40). Each modality is reduced to 50 components *Z*^(*m*)^, and a mutual 50-nearest-neighbour graph is built and symmetrically normalised. Its eigenvectors Φ^(*m*)^ and eigenvalues **Λ***_m_* give multiscale diffusion coordinates at three diffusion times,

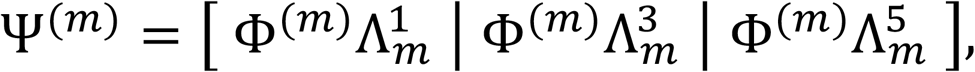

where raising **Λ***_m_* to the power *t* down-weights rapidly decaying directions, so shorter diffusion times preserve local structure while longer times capture broader structure. Three costs follow, comparing manifold position, reduced features and the shared-feature profiles *H*^(*m*)^:

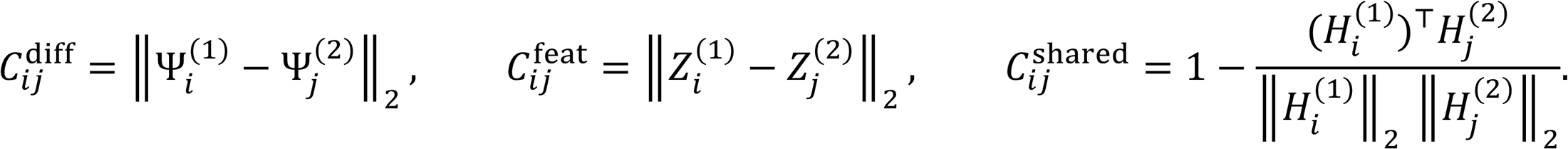

Each cost is scaled to the interval [0,1], written *C*^?^, and the three are combined as

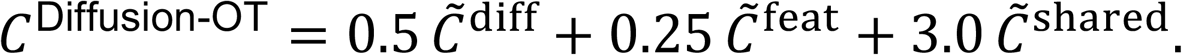

The shared-feature term receives the largest weight so that manifold geometry remains anchored to direct cross-modal evidence.

CCA-OT combines direct shared-feature similarity with anchor-based canonical correlation analysis. The selected shared features are z-scored within each modality, and each cell is normalised to unit length, giving 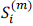 and the direct cost 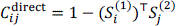. Separately, reciprocal cross-modal neighbours define an anchor set *A*, where (*i*, *j*) ∈ *A* only if *j* is among the 15 nearest neighbours of *i* and *i* is among the 15 nearest neighbours of *j*. Canonical correlation analysis (17) is fitted to those anchored rows alone,

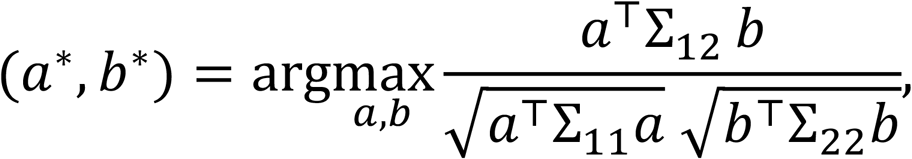

in which Σ_11_ and Σ_22_ are the within-modality covariance matrices and Σ_12_ is the cross-covariance between anchored rows. All cells are then projected into the canonical space as *U* and *V*, giving a second cost 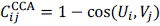. The two costs are fused by row rank, and the fused rank matrix *Q* becomes a coupling through reciprocal rank,

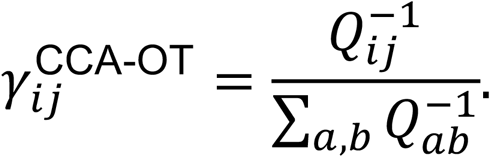

Ensemble-OT converts each component to row ranks, so that rank 1 always means the best partner, giving *R^D^* from Diffusion-OT, *R^C^* from CCA-OT and *R^M^* from MaxFuse. The fused cost is

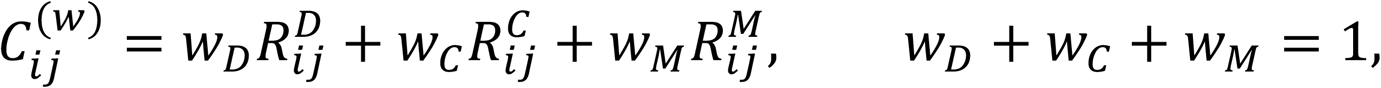

searched over the simplex in steps of 0.025. The weights are chosen without the true pairing and without any cell-type label. The similarity *S* = −*C*^(w)^ is corrected for hubness by cross-domain similarity local scaling,

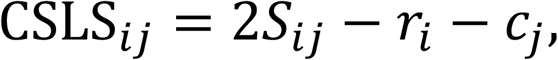

where *r_i_* and *c*_j_ are the mean similarity of cell *i* and of cell *j* to their selected nearest cross-modal neighbours. A pair therefore receives a high score when the two cells are relatively specific partners rather than broadly like many cells. The chosen weights maximise the average CSLS value over mutually best-matching pairs (59).

Entropic transport. The costs from Diffusion-OT, Ensemble-OT and MaxFuse become couplings through Sinkhorn balancing (11). The kernel is

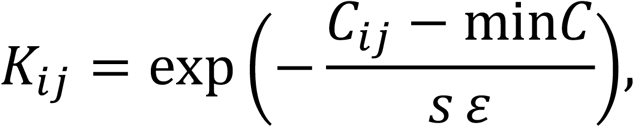

where *ε* controls the concentration of the transport plan, with smaller values producing more concentrated couplings and larger values producing more diffuse couplings, and *s* is a scale factor that is either one or the median cost. Two scaling vectors are updated in turn, *u* ← *a* ⊘ *Kv* and *v* ← *b* ⊘ *K*^T^*u*, until the row and column masses match the uniform marginals *a_i_* = 1/*n*_1_ and *b*_j_ = 1/*n*_2_. The coupling is then

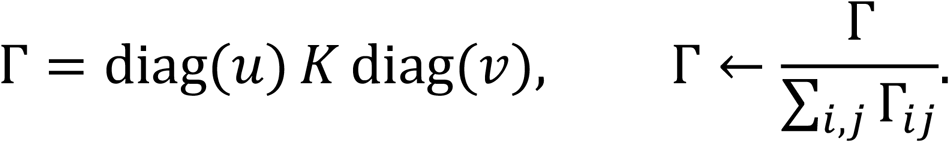

Uniform marginals prevent a small number of cells from absorbing most of the correspondence mass, and the final rescaling gives unit total mass. CCA-OT instead fuses its Sinkhorn-derived components by within-row rank, leaving column masses unconstrained; downstream analyses renormalise each row and therefore use only row-wise mass.

### Ambiguity detection and cell selection

We define ambiguity as uncertainty in the cross-modal correspondence of one cell, rather than as an intrinsic property of that cell, and a larger score always means a less resolved match. One score serves all three variants, which differ only in where the reference groups come from.

For a query cell *i*, the coupling row is first turned into a distribution over reference cells,

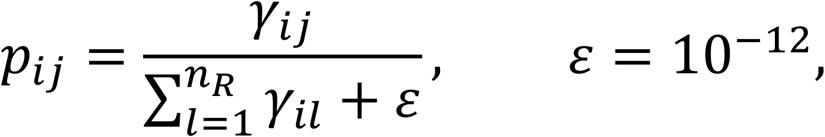

where *γ_i_*_j_ is the correspondence mass from query cell *i* to reference cell *j* and *n_R_* is the number of reference cells. Every reference cell *j* carries a group label *g*_j_, so the mass is pooled by group and renormalised over the *K* groups,

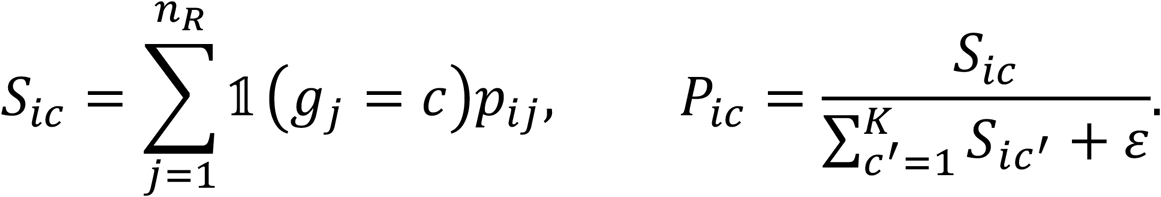

Writing *P_i_*(_1_) and *P_i_*(_2_) for the largest and second-largest votes, the score is

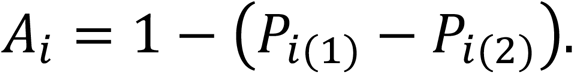

A large gap means one group clearly wins and ambiguity is low, whereas a near tie means the evidence is split between two interpretations. RefA-Amb takes the groups from the annotated cell types of modality 1, RefB-Amb from those of modality 2, and LF-Amb from Leiden clusters of the reference modality, so LF-Amb needs no labels at all (60).

These scores rank cells but do not say how many to flag. Because the sorted curve is close to a straight line, each score is first transformed by *ã_i_* = exp(10(*a_i_* − 1)), which is strictly increasing and therefore leaves the ranking untouched while making the bend visible. The transformed scores are sorted from largest to smallest, and the L-method fits two least-squares lines and takes the split that minimises the size-weighted error

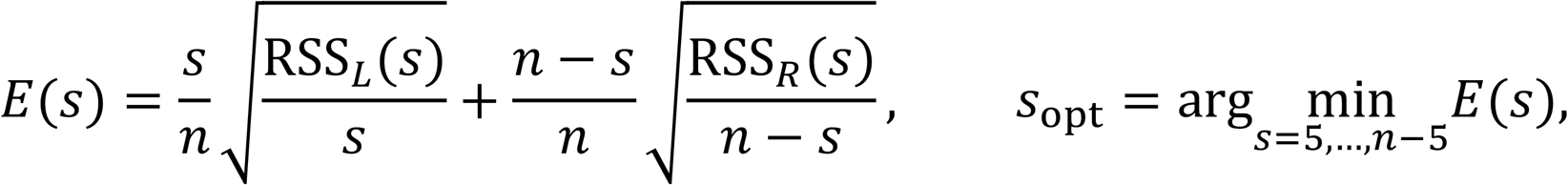

in which RSS*_L_* and RSS*_R_* are the residual sums of squares of the two fitted lines and the two weights stop a very short segment from winning only because it is short. The number of flagged cells is *k* = min(*s*_opt_, ⌊0.5*n*⌋), so no more than half of a dataset can be removed, and every run in which the cap binds is recorded.

### Hyper-parameter tuning

All external tools were tuned with one protocol. For each modality pair we used two small datasets and, where one existed, an intermediate dataset, and kept the configuration with the best mean score. Small grids were searched fully and large grids by greedy coordinate search from each tool’s documented default, which keeps the run count affordable while still moving every parameter. Correspondence methods were scored by symmetric FOSCTTM, which needs a known pairing and was therefore computed on co-assay pairs only, over 1,563 runs. Label-transfer tools were scored by macro F1 over 1,326 runs, and integration tools by 0.6 times NMI plus 0.4 times batch mixing over 1,392 runs (supplementary method).

Two kinds of parameter mattered most, and the selected values varied by modality pair. For the correspondence tools these were the entropic regularisation and the number of features kept: SCOT chose k of 30 to 50 with epsilon between 0.001 and 0.01; UnionCom settled on kmax 5, epsilon 0.001 and beta 0.5 in both groups it completed; MMD-MA chose a latent dimension of 10 or 30 with an orthogonality weight of 0.01 or 0.1 and a distortion weight of 1 or 10; and MaxFuse chose 10 to 20 neighbours, 15 components throughout, and a blending weight of 0.5 except on RNA-methylome, where it fell to 0.3. For the integration and label-transfer tools these were neighbourhood size and dimensionality. In integration, Seurat chose dims 10 to 30 with k_anchor 5 to 30 and k_weight 20 to 100; Symphony dims 10 to 30 with 500 to 5,000 variable genes; Harmony theta 2 to 8 with lambda 0.1 to 5; fastMNN k 10 to 50 with d of 20 or 50; Scanorama knn 5 to 50 with sigma 5 to 100; scVI 20 to 100 latent dimensions with one to three layers; and WNN, which runs on co-assay pairs only, knn 5 to 50 with 10 to 20 RNA and 18 to 25 protein dimensions. In label transfer, Seurat chose dims 20 to 75 with k_anchor 10 to 30 and k_weight 10 to 50; Symphony knn_k 5 to 50 with dims 20 to 100; Harmony k 5 to 30 with 30 to 100 components; SingleR the Wilcoxon marker rule in four of the six groups with 100 or 200 markers; scmap 200 to 3,000 features with the assignment threshold at zero everywhere except co-assay RNA-ADT, where it was 0.7; ItClust a learning rate of 0.0005 to 0.005 with 50 to 200 epochs; and Concerto a learning rate of 0.0001 in every group with 5 to 40 pre-training epochs. Settings were selected for every modality pair and carried unchanged to the diagonal pairs of the same type.

The models developed here were tuned on the same datasets, with the same metrics and the same search. Among the couplings, Diffusion-OT chose 3,000 variable features on RNA-ADT and 20,000 on the other two groups, with epsilon 0.05 on RNA-ADT and RNA-ATAC and 0.005 on RNA-methylome, and a diffusion weight of 0.75 on RNA-ADT and 0.25 elsewhere. Diffusion-GW kept epsilon at 0.005 and three refinement rounds in every group, with 3,000 or 20,000 features and a geometry weight of 0.5 or 1.0. CCA-OT was the most stable, choosing 2,000 shared features, 30 canonical dimensions and 15 anchor neighbours in all three groups, with the transport epsilon at 0.05 on RNA-ADT and 0.1 on the other two. Ensemble-OT has one parameter, the CSLS neighbourhood, which the sweep set to 10 on RNA-ADT and RNA-methylome and 5 on RNA-ATAC. For integration, AnchorAE trained for 80 to 300 epochs with a mutual-neighbour weight of 0.1 to 1 and 5 to 15 neighbours, AnchorFormer for 80 to 200 epochs with a weight of 0.1 to 2 over the same neighbour range, and AnchorMix used a 16-to-64-dimensional embedding trained for 100 to 400 epochs at a learning rate of 0.0005 to 0.005. For label transfer, the two nearest-neighbour heads tuned a single parameter, the number of neighbours, which ran from 3 to 30 for AnchorAE and from 3 to 50 for AnchorFormer, and AnchorAE-HGB chose a learning rate of 0.05 or 0.1 with 100 to 800 boosting iterations.

### Custom data integration and label transfer models

We developed three integration models. AnchorAE and AnchorFormer share one input, one latent size and one training objective, and differ only in the encoder body. AnchorMix is a separate design with one encoder per modality.

AnchorAE is an autoencoder trained on the two modalities at once (61, 62). The input is the shared-gene matrix, z-scored within each modality so that the two blocks sit on a common scale without changing the structure inside either. The encoder maps the *F* input features to 256 hidden units through a linear layer, layer normalisation and a GELU activation, and then to a 50-dimensional latent code *z*; the decoder mirrors this back to *F*. Training minimises

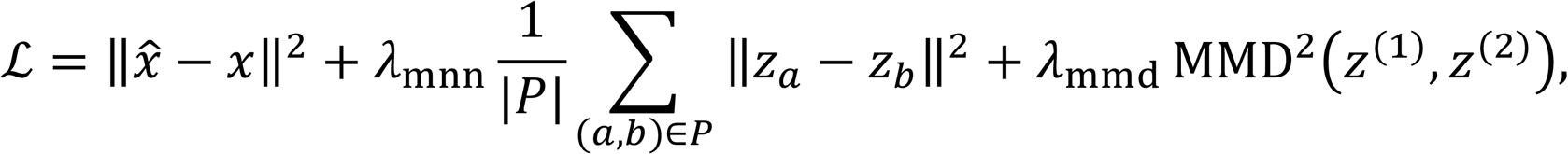

where the first term is reconstruction, *P* is the set of mutual nearest-neighbour pairs found across the two modalities in the current latent space, and the last term is a radial-basis maximum mean discrepancy between the two modality distributions, summed over bandwidths 0.5, 1 and 2. The pairs in *P* are recomputed every 15 epochs after a 20-epoch warm-up, so alignment follows the embedding as it forms. Only reciprocal pairs are used, which pulls shared cell types together while leaving populations present in one modality alone.

AnchorFormer uses the same input, latent size, decoder and objective, and inserts three self-attention blocks between the input layer and the latent projection (63). Each block updates its hidden state as

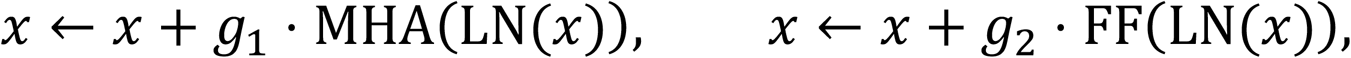

where MHA is eight-head self-attention across cells, FF is a two-layer feed-forward network, LN is layer normalisation, and *g*_1_ and *g*_2_ are learnable scalar gates initialised at 0.1. The gates are the point of the design. Attention across cells can mix cell identities and blur cell-type structure, so starting the gates near zero makes the block behave almost like the identity at the beginning of training, and attention is used only where it lowers the loss. AnchorFormer can therefore refine the representation that AnchorAE would learn, but cannot easily destroy it.

AnchorMix encodes each modality with its own network into one shared embedding, and adds guarded mixing terms on top of reconstruction. A fixed anchor is computed once from a joint Laplacian in principal-component space and is never rebuilt during training, because rebuilding it from the current embedding was unstable. Unsupervised pseudo-clusters are taken from that anchor by k-means, using no cell-type labels. The objective is a weighted sum,

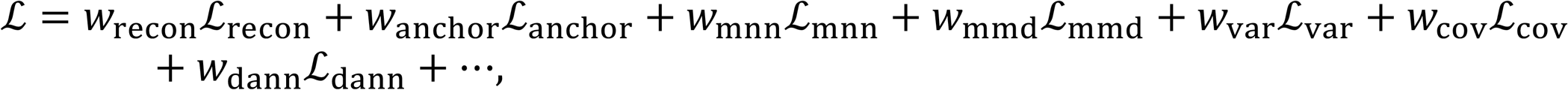

in which ℒ_anchor_ ties the embedding to the fixed anchor, ℒ_mnn_ attracts reciprocal cross-modal pairs, ℒ_mmd_ matches the two modality distributions inside each pseudo-cluster rather than globally, ℒ_var_ and ℒ_cov_ keep each latent dimension informative and uncorrelated, and ℒ_dann_ is a class-balanced adversarial term that makes the modality hard to predict from the embedding. Further terms pull cells toward their own pseudo-cluster centroid and attract high-confidence reciprocal coupling pairs that fall in the same pseudo-cluster. Every mixing term is conditioned on a pseudo-cluster, so the model mixes modalities within a cell population instead of across the whole dataset, which is what protects cell-type structure.

Three custom label-transfer models are reported, each an encoder paired with a simple classifier applied to the 50-dimensional embedding that encoder produces, so the model compared is the pair rather than the classifier alone. These embeddings come from the same encoders evaluated in the integration benchmark. AnchorAE applies k-nearest neighbours to the AnchorAE embedding, giving a query cell the majority label of its k closest reference cells, and AnchorFormer applies the same classifier to the AnchorFormer embedding; for both, k is the only tuned parameter. AnchorAE-HGB applies a histogram gradient-boosting classifier to the AnchorAE embedding and tunes four parameters: the number of boosting iterations, the learning rate, the maximum leaves per tree, and the L2 penalty on the leaf values.

### DI-ElasticFlow coupling upgrade

ElasticFlow is a training-free refinement applied after any integration method, so Seurat, Harmony, fastMNN and the custom encoders are never refitted. Reference coordinates stay fixed and only the query block moves. The coupling says where each query cell should go, and a graph-Laplacian penalty keeps neighbouring query cells moving together. Let the baseline embedding and the coupling be

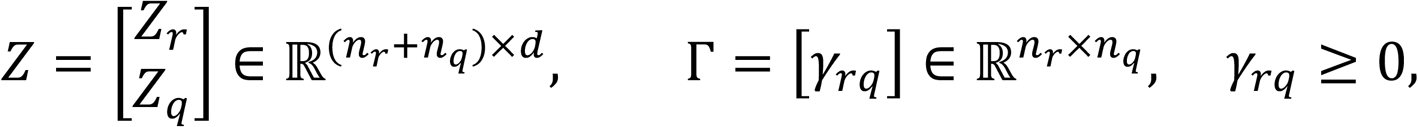

where the upper block holds the fixed reference coordinates, the lower block the movable query coordinates, *d* is the embedding dimension, and *γ_rq_* is the correspondence mass between one reference cell and one query cell. For each query cell *q*, ElasticFlow keeps the *k*_coup_ reference cells with the largest coupling values, renormalises their weights to sum to one, and takes their weighted barycentre as the preferred position,

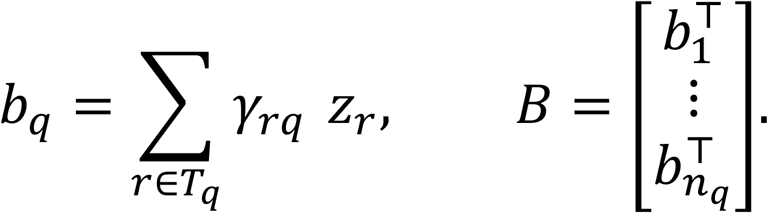

The barycentre is the point that minimises the weighted squared distance to those partners. Moving a cell onto it outright is avoided, because uncertain partners would collapse the query manifold onto the reference. A *k*_smooth_-nearest-neighbour graph is therefore built among the original query coordinates and symmetrised, giving the degree matrix and the combinatorial Laplacian

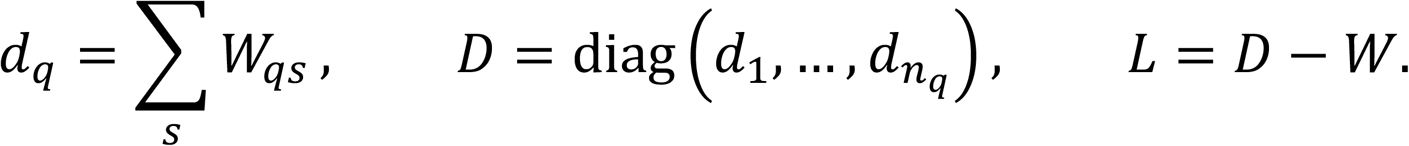

Let *U* hold one displacement vector per query cell. ElasticFlow balances attraction to the targets against a roughness penalty that is small when neighbouring cells move alike. The objective is quadratic, so its stationary point is one sparse symmetric positive-definite system,

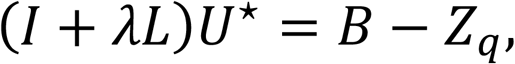

in which *λ* sets the stiffness: a small value lets each cell follow its own target, and a large value forces neighbouring displacements to agree. A second parameter controls how much of the displacement is applied,

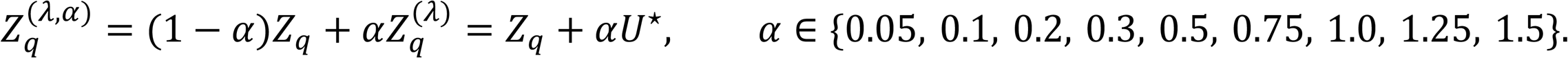

A candidate is accepted only when three label-free conditions hold together, a test we refer to as Safe-upgrade rule: the integration LISI score rises above its baseline value, at least half of each query cell’s original fifteen nearest neighbours survive the correction, and the structure proxy does not fall below its baseline value,

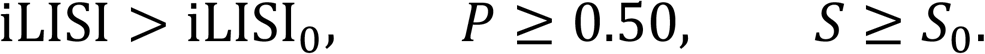

Writing *θ* = r*α*, *λ*, *k*_coup_, *k*_smooth_s for one configuration and *A* for the set of accepted configurations, ElasticFlow applies

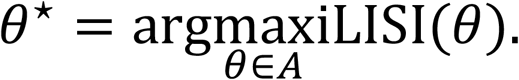

If *A* is empty, the baseline embedding is returned exactly unchanged. This fallback is the do-no-harm guard inside Safe-upgrade: an embedding may be replaced only by one that mixes the two modalities better while keeping local neighbourhoods and cluster structure at least as good as before. Nothing in the rule uses cell-type labels, and NMI, ARI and the two silhouettes are recorded for every candidate but never decide which candidate wins. The four parameters are tuned separately for every dataset, tool and coupling by two passes of coordinate descent, which cost 80 evaluations against a full grid of 9,360 combinations.

WNN needs separate treatment, because it returns a weighted shared-nearest-neighbour graph with per-cell modality weights rather than coordinates (23). ElasticFlow acts on the embedding derived from that graph and leaves the graph itself untouched, so a gain measured there should not be read as a change to WNN’s own output. WNN is also restricted to the 29 co-assay pairs, because a per-cell modality weight is undefined without measured cell pairing.

Only the query coordinates are replaced, so the final output is

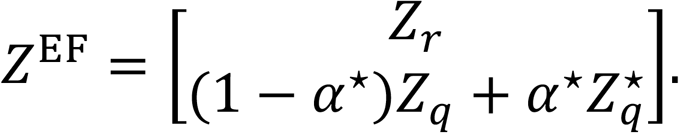

ElasticFlow was applied independently with the MaxFuse, Ensemble-OT, CCA-OT and Diffusion-OT correspondences.

### LT-BlendGate coupling upgrade

Each label-transfer run had four comparable arms: the unmodified baseline tool, the LT-BlendGate prediction with all query cells retained, the upgraded prediction after ambiguity-guided removal of query cells, and the upgraded prediction after count-matched random removal. The two transfer directions were evaluated independently. LT-BlendGate adds the coupling to a baseline prediction without retraining the tool and without touching its embedding, so the coupling acts as an independent second source of evidence. Let *Y* ∈ {0,1}*^nR^*^×*K*^ be the one-hot matrix of reference labels over *K* cell types and let Γ be the coupling. Because the coupling links cells rather than classes, its mass is pooled over the reference cells of each type,

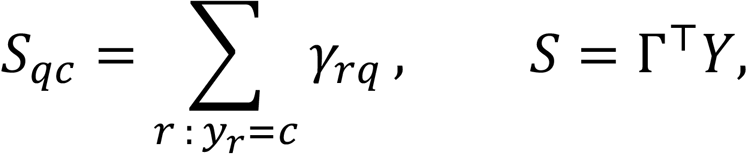

so that several moderate links to one cell type can together give strong evidence even when no single reference cell dominates. The pooled scores are clipped at zero, divided by each cell’s largest value, and then sharpened,

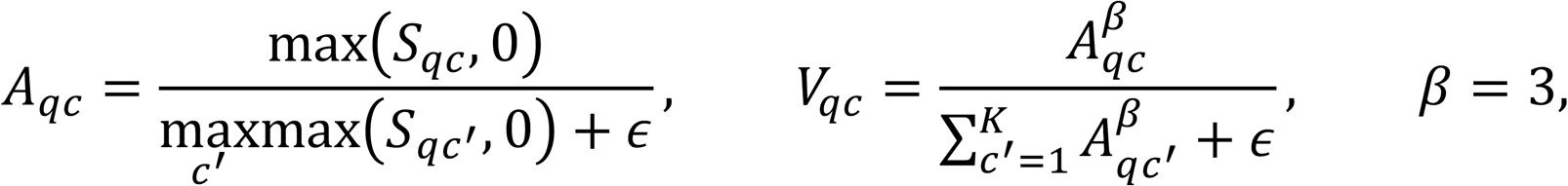

where the exponent *β* makes a clear preference clearer while leaving a flat vote flat, and *ε* = 10^−9^ prevents division by zero. The baseline method supplies its own class distribution 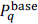. Two scalars then decide how much the coupling may contribute: the baseline confidence 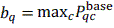 and the vote sharpness 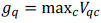. The blend weight is their product,

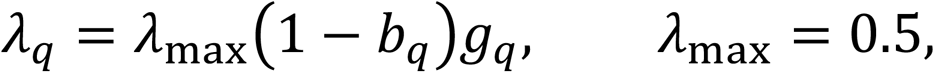

which behaves as an AND gate. The coupling gains influence only when the baseline is unsure and the vote has a clear preference, so a confident baseline is left alone and a flat vote cannot overturn one. The final score is the convex blend

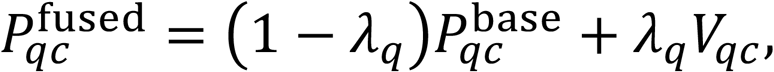

and the upgraded label is the class with the largest fused score. The procedure runs independently in both transfer directions, using Γ^T^ for the reverse direction and recomputing every quantity, and was evaluated separately with the MaxFuse, Ensemble-OT, CCA-OT and Diffusion-OT correspondences. Ambiguity-guided removal is applied only after the full-reference prediction has been computed.

## Acknowledgements

During the preparation of this manuscript, the authors used ChatGPT (OpenAI) and Claude (Anthropic) to assist with language editing, improving clarity and organisation, and refining text written by the authors. All AI-assisted suggestions were critically reviewed and edited by the authors.

## Supplementary files

Datasets_composition_supplementary.pdf

Methods_supplementary.pdf

Supplementary figures.pdf

Ambiguity_elbow_plots.zip

Label_transfer_plots.zip

Label_transfer_supplementary.xlsx

Data_integration_plots.zip

Label_transfer_supplementary.xlsx

